# Modeling nascent transcription from chromatin landscape and structure

**DOI:** 10.1101/2024.06.04.597340

**Authors:** Marc Pielies Avellí, Arnór Sigurdsson, Takeo Narita, Chunaram Choudhary, Simon Rasmussen

**Affiliations:** Novo Nordisk Foundation Center for Protein Research, Faculty of Health and Medical Sciences, University of Copenhagen, Copenhagen N, Denmark; Novo Nordisk Foundation Center for Basic Metabolic Research, Faculty of Health and Medical Sciences, University of Copenhagen, Copenhagen N, Denmark; Novo Nordisk Foundation Center for Genomic Mechanisms of Disease at the Broad Institute of MIT and Harvard, 415 Main St. Cambridge, MA 02142, USA

**Keywords:** EU-seq, nascent transcription, chromatin landscape, Micro-C, deep neural networks

## Abstract

**Background:** Different cell types and their associated functionalities emerge from a single genomic sequence when certain regions are expressed while others remain silenced. Modeling gene expression and its potential malfunctioning in different cellular contexts is hence pivotal to understand both development and disease.

**Results:** We present the Chromatin Landscape and Structure to Expression Regressor (CLASTER), an epigenetic-based deep neural network that can integrate different data modalities describing the chromatin landscape and its 3D structure. CLASTER effectively translates them into nascent transcription levels measured by EU-seq at a kilobasepair resolution. Our predictions reached a Pearson correlation with targets above r=0.86 at both bin and gene levels, without relying on DNA sequence nor explicitly extracted chromatin features. The model mostly used the information found within 10 kbp of the predicted locus, even when a wide genomic region of 1 Mbp was available. Explicit modeling of long-range interactions using multi-headed attention and high-resolution chromatin contact maps had little impact on model performance, despite the model correctly identifying elements in these inputs influencing nascent transcription. The trained model served then as a platform to predict the transcriptional impact of simulated epigenetic silencing perturbations.

**Conclusions:** Our results point towards a rather local, integrative and combinatorial paradigm of gene regulation, where changes in the chromatin environment surrounding a gene shape its context-specific transcription. We conclude that the predominant locality and limitations of current machine learning approaches might emerge as a genuine signature of genomic organization, having broad implications for future modeling approaches.

## Background

A single genome gives rise to a wide variety of cells, with diverse morphological properties and functions. These features of the cells emerge when certain regions of the DNA are expressed while others are kept silent, a process known as gene regulation. The rules governing such regulatory processes in different cellular contexts still constitute a highly debated topic, and their understanding is crucial to study development and disease. Many models of gene regulation have already been proposed approaching the problem at different biological layers [1–5]. At a high level, gene expression is modulated by the interplay of a set of regulatory elements, e.g. enhancers and promoters, which have been shown to be broadly compatible in most cases [6,7]. Their activities are usually described using a number of epigenetic marks and can be combined multiplicatively to shape RNA transcription [7–9]. The establishment of functional pairings between these regulatory elements is then conditioned by their contact frequency, given by the three-dimensional structure of chromatin. Models such as the Activity by Contact (ABC) [10] and the more recent ENCODE-rE2G [11] exploit these features.

Regulatory elements and their associated genes are ultimately encoded in the DNA. Many computational approaches have therefore put a special emphasis on the genomic sequence, identifying patterns and motifs, integrating them and predicting a set of output tracks describing various downstream genomic and epigenomic processes [1–5]. The advent of large language models (LLMs) is also powering a new wave of sequence-based machine learning models such as Nucleotide Transformer [12], Borzoi [13], DNABERT [14], HyenaDNA [15] or Evo [16]. These present improvements in several fronts, e.g., being more robust to genomic variability, extending the studied genomic regions or achieving higher resolutions while bringing down the computational costs. Finally, new advances in the 3D organization field and chromatin conformation capture techniques have brought to light finer structural details (Micro-C) [17,18] and also show promise to improve predictive modeling [19,20].

However, this explosion in the genomics analysis toolbox is mostly inherited from the computer vision and natural language processing fields, which pursue different objectives and describe data modalities that attend to highly different sets of rules. This is best illustrated by the fact that even some of the most advanced machine learning models rely primarily on the close neighborhood of the predicted points, without paying attention to most of the sequence provided [9]. This is of particular interest in the case of attention-based deep learning models, which handle the complete sequences at once and therefore do not impose any spatial constraints. This raises the question: are these machine learning models predominantly local because of current technical limitations and training schemes, or is this a property of the underlying data distribution, i.e., a signature of genomic organization induced by elements such as evolutionary pressure?

Here we present Chromatin Landscape and Structure to Expression Regressor (CLASTER), a deep convolutional neural network aimed to learn the mapping from a given chromatin landscape and its three-dimensional structure to the corresponding nascent RNA transcription profiles. CLASTER constitutes a cell-type agnostic model that integrates properties of both chromatin state and structure without relying on the genomic sequence. The model can identify patterns between chromatin marks and across the genome at several levels of abstraction and combine them in a non-linear fashion. Of note, the multiplicative nature of enhancer and promoter activities deeply motivates the use of non-linearities, which would not be captured by linear models [1]. Similar principles have already been successfully applied in gene expression classification [21,22] and regression [19], and contact map prediction [23,24]. Motivated by recent findings of broad compatibility between regulatory elements, we identified the core set of elements required to predict the nascent transcription landscape from an epigenetic perspective. The trained models were then used to explore the power of new perturbational approaches on such data to simulate the transcriptional impact of epigenetic remodeling, which has already shown great therapeutic potential [25].

## Results

### CLASTER integrated multi-modal chromatin data to predict nascent RNA levels in a sequence-agnostic manner

We developed CLASTER, a deep neural network designed using the EIR framework [26] and trained it to translate a given chromatin landscape and its corresponding high-resolution 3D contact frequency map to the matching target nascent RNA profiles (**Figure 1a, Supplementary Figure 1, Supplementary Table 1**). Nascent transcription levels were obtained for mouse Embryonic Stem Cells (mESCs) from Narita et al. 2021 [27], where they had been measured by RNA labeling with 5-ethynyl uridine and sequencing (EU-seq). CLASTER was trained to predict kilobasepair resolution EU-seq profiles in a wide region, spanning from -200 kbp to 200 kbp from the TSS of a gene of interest that was located at the center of the predicted window (**Figure 1b, Supplementary Figure 2, Methods**). Chromosomes 17 and 4 were used to obtain our validation and test set regions, respectively (**Supplementary Table 2**).

**Fig 1.**
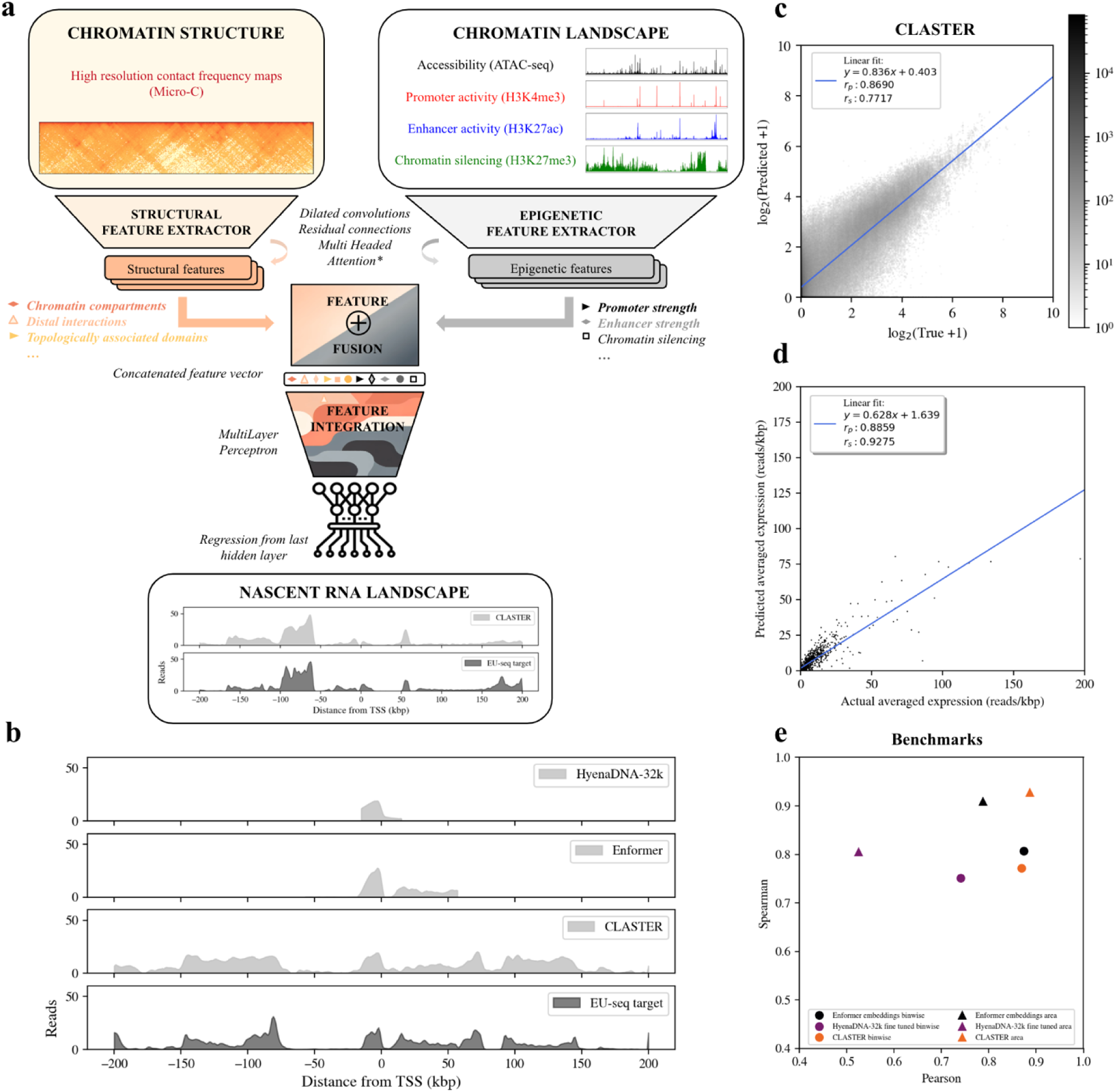
CLASTER can translate the chromatin landscape and structure to the corresponding nascent RNA profiles in mESCs. **a)** Model schema. CLASTER maps the epigenetic landscape of a chromatin region of 1Mbp and its matching contact frequency map to the corresponding nascent expression levels in a window of 401 kbp. **b)** Targets and predictions around the gene Tmem222 (chr4) for the different models. Prediction ranges were adapted to the context length of each model. **c)** Density plot showing log-transformed, binwise predictions vs. target values in all test samples, i.e. across protein coding genes in chromosome 4. **d)** Length normalized nascent transcription levels for the central gene in each predicted sample. Normalized transcription levels were obtained by integrating the area under the EU-seq curve inside the gene boundaries and normalizing by gene length. **e)** Pearson and Spearman correlations between test predictions and targets at binwise and gene levels for the different models.

To perform the predictions, the network used two matrices representing the different data modalities, which were processed in separate branches of the model. The first branch processed four complementary genomic tracks that characterized the epigenetic landscape in a chromatin region of 1Mbp at a 100bp resolution. These four core tracks describe chromatin accessibility (ATAC-seq), promoter activity (H3K4me3), promoter and enhancer activity (H3K27ac) and heterochromatin (H3K27me3), respectively. An optional second branch of the model was aimed at processing high resolution structural data. This was given in the form of Micro-C contact frequency maps exploring also 1Mbp of genomic context at a 1.6 kbp resolution. These maps described contact frequencies between different parts of the genomic region explored in each sample and were obtained for the same cell line from [28]. The model then extracted features in both data modalities separately using a set of dilated convolutions and residual connections (**Figure 1a, Supplementary Figure 1**). Optional multi-headed attention layers were then added to the landscape branch to ease the processing of the sequential appearance of such features and long-range interactions between them. The resulting high-level features were then combined and integrated using a set of fully connected layers, and finally a dense layer connected the last hidden layer to each output separately.

CLASTER displayed a high test prediction accuracy, with Spearman and Pearson correlations between log transformed predicted values and ground truth signal intensities of r_s_=0.7717 and r_p_=0.8690 across all predicted bins (**Figure 1c)**. Test performances increased to r_s_=0.9275 and r_p_=0.8859 when comparing the expected nascent transcription levels inside target gene boundaries (**Figure 1d, Methods)**, where a lower number of near zero values would be expected. In summary, CLASTER predicted nascent RNA levels given the corresponding chromatin landscape and its matching high-resolution contact frequency map. This was achieved by extracting features at different levels of abstraction from both data modalities without relying on the DNA sequence. Of note, these predictions comprised most protein coding genes in chromosome 4, with no filters on specific ranges of expression levels nor their enhancer-dependence status.

### Nascent transcription can be predicted from DNA sequence

The enrichment levels of different chromatin marks in a given locus, which define the chromatin states [29], may be interpreted as epigenetic embeddings containing relevant functional information of that genomic region. This makes them somewhat similar in role to the inner sequence representations of DNA-based models like the Enformer [2] or HyenaDNA [15], and therefore we wanted to compare their respective capabilities to predict the nascent RNA landscape described by EU-seq.

Sequence embeddings of the central 160kbp regions matching our samples were obtained for the Enformer model with pretrained, frozen weights (**Supplementary Figure 3**, **Methods**). We then trained a simple head mapping the embeddings to the target EU-seq outputs (**Supplementary Figures 4-5, Methods**). The performance of the model head on Enformer’s pretrained embeddings, i.e., with no fine-tuning nor further training required, achieved Pearson and Spearman correlations of r_s_=0.8067 and r_p_=0.8739 for the test predictions (**Figure 1e, Supplementary Figures 6-8**). However, the target window had to be shortened given Enformer’s narrower context length and cropping of the latent sequence representations **(Figure 1b, Supplementary Figure 8**). Sequence embeddings obtained by pretraining the DNA language models HyenaDNA-160-kbp and 32-kbp in a next nucleotide prediction task could not be used in the same manner to directly predict nascent expression levels in a regression fashion (**Methods**). The embeddings could however be tailored to our task by fine-tuning the pretrained backbone and the added head together. Predictions using a fine-tuned HyenaDNA-32k model displayed Pearson and Spearman correlation coefficients of r_s_=0.7514 and r_p_=0.7403 (**Figure 1e, Supplementary Figure 6**).

In conclusion, DNA-sequence based models trained in a supervised manner like the Enformer provide embeddings encoding valuable functional information, which can be directly probed to predict nascent transcription levels. These results were to be expected since the Enformer was originally trained to predict 1,643 mouse genomic tracks, including RNAPII binding and a number of CAGE experiments that were highly related to our outputs. HyenaDNA, a genomic sequence foundation model, could also be used to predict the nascent transcription landscape. However, sequence embeddings obtained from pretraining HyenaDNA in a next nucleotide prediction task did not encode the information necessary to predict our targets, requiring further fine-tuning of both HyenaDNA backbone and the added model head.

### CLASTER learned the functional roles of chromatin states rather than those of specific marks

*In silico* perturbations of the inputs can allow us to retrieve some of the information learned by the models, while enabling the simulation of arbitrary epigenetic variability. They might be used, for instance, to identify potential transcriptional changes induced by an epigenetic silencing of chromatin, a valuable alternative to genetic knock-out [25]. To this end, we performed virtual perturbational experiments aimed to simulate a loss of the activity of single active enhancers using only the chromatin landscape branch of CLASTER (**Methods**). Proximal and distal Enhancer Like Signatures (pELS, dELS), were obtained from [30] (**Supplementary Table 3**).

We found that the enrichment levels of different chromatin marks in native chromatin were entangled, i.e., they did not explore all possible values but were only found in a set of allowed combinations encoding chromatin states with diverse functional roles [29] (**Figure 2a**). Perturbing the enhancer mark H3K27ac alone (P1) had a modest impact on the predicted profiles. An isolated *in silico* ablation of H3K27ac levels in active enhancer loci yielded still active chromatin states with unaltered accessibility and background H3K4me3 levels, unlike what would be expected *in vivo* [27] (**Figure 2b, top)**. The integrative nature of the convolutions across tracks in the inputs led the model to map chromatin mark combinations to the corresponding RNA landscape instead of assigning independent roles to different input tracks. This limited the ability of single mark perturbations to infer biologically meaningful phenotypes, but illustrated that the ratios between marks and their combined contributions rather than isolated mark enrichments define the regulatory role of a genomic region [31,32].

**Figure 2.**
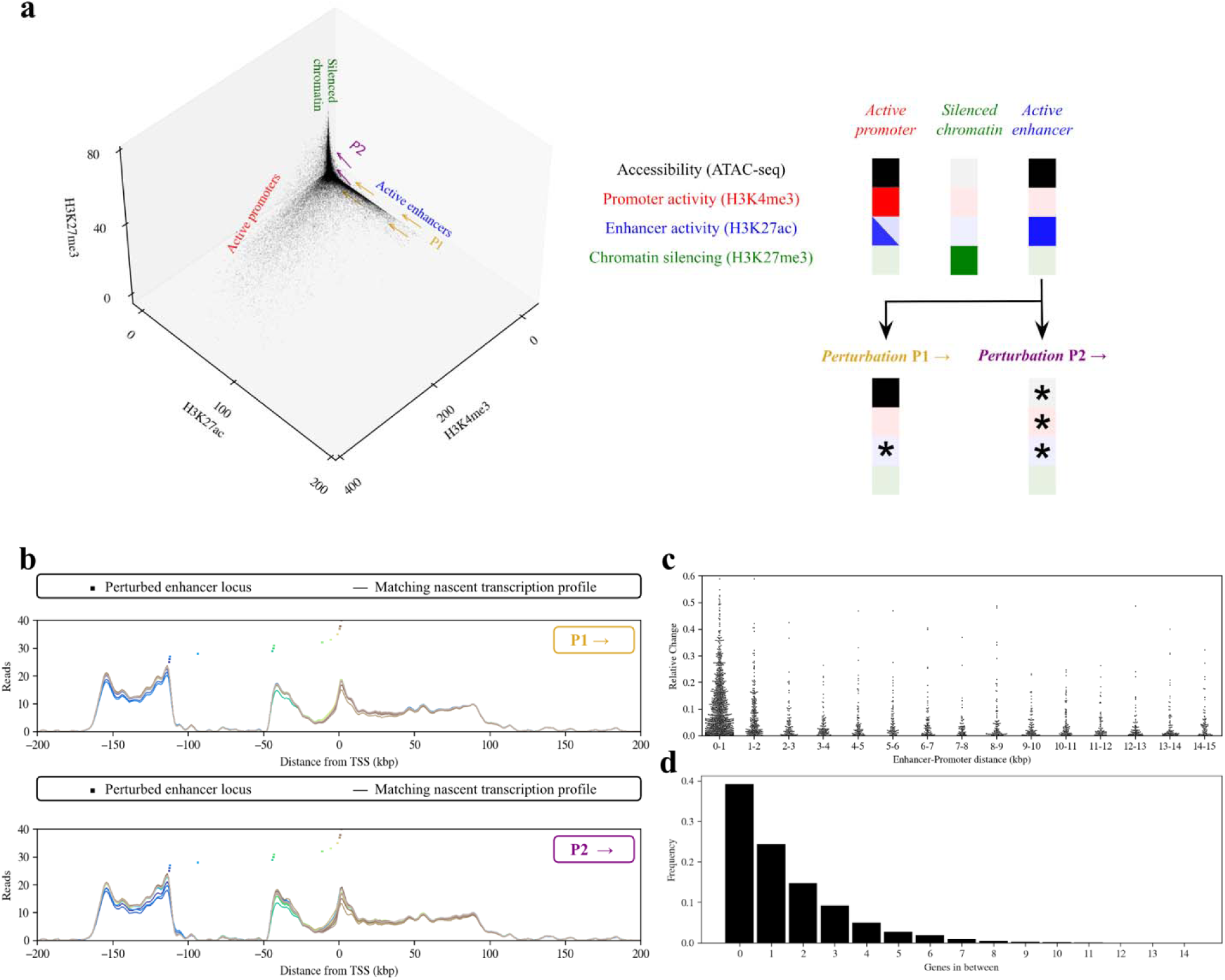
Local silencing of enhancer loci mainly affects the neighboring genes. **a)** Combinations of chromatin mark enrichment values at a given bin sampled from chr 4. Example chromatin states and the resulting changes induced by perturbations P1 and P2 are illustrated. **b)** Predicted output EU-seq profiles from inputs where only the enhancer mark H3K27ac has been ablated at a given enhancer locus (P1) and inputs with silenced chromatin state substitution at the enhancer locus (P2). The ablated enhancer loci, shown as small rectangles, and their corresponding output profiles are depicted using the same color. Shown predictions are centered at the protein coding gene Tmeff1 (ENSMUSG00000028347.14, chr 4). **c)** Relative change in length normalized area under the EU-seq curve for each gene when silencing an enhancer (P2) at a given distance of the gene’s TSS. Shown are all enhancer-promoter pairs left after preprocessing of test samples. **d)** Number of cases in which the enhancer ablation affected the most a gene located on either side (adjacent) vs the enhancer skipped a number of genes. Results are shown for P2.

### *In silico* silencing of enhancer loci induces mainly the downregulation of close genes

To better simulate the effects of an epigenetically silenced enhancer, the whole chromatin state at each target enhancer locus was substituted for a heterochromatic-like state (P2), enhancer by enhancer. An arbitrary silenced state was obtained by setting all marks except the silencing H3K27me3 to zero reads (**Methods**). The introduced chromatin state substitutions mainly modified the predicted expression profiles around the perturbed point in a distance-dependent fashion, having an influence on a varying number of genes (**Figure 2b, bottom**). Interestingly, the perturbations led to a sustained decrease in EU-seq across the body of the affected domains while keeping the overall structure characterizing the profiles. Clusters of neighboring ELS affected the same region, pointing towards a coordinated action of several enhancers [11]. Furthermore, the degree of downregulation that could be exerted by *in silico* silencing an active enhancer on any given gene presented a clear inverse relation with the distance separating them, decaying rapidly after a few kilobasepairs (**Figure 2c, Supplementary Table 4**). From a complementary point of view, the enhancers that had the highest impact on a gene’s nascent transcription levels were in a high fraction adjacent to the gene (ca. 40%), with less and less significant cases skipping increasing numbers of genes (**Figure 2d, Supplementary Figure 9**).

To summarize, predicting the EU-seq levels in a genomic region of 401 kbp per sample allowed us to assess the range and degree of influence of a set of tailored *in silico* perturbations in enhancer loci. The effects of such perturbations were not propagated towards regions far from the perturbed locus, indicating a predominantly local behavior of the network. CLASTER linked the perturbations in those loci to the adjacent domains with significant EU-seq enrichment, modifying in a distance-dependent manner the transcription profiles and therefore affecting a varying number of genes found around each perturbed locus.

### Most of the provided context was not used for the predictions

To disentangle the contributions of all marks, we used the integrated gradients attribution method [33]. This axiomatic attribution mechanism enabled us to quantify the relative importance of the different chromatin marks at all measured loci towards the predicted EU-seq levels at each target locus (**Figure 3a**). Positive and negative attribution scores indicate direct or inverse associations between input and output enrichment levels, respectively. The resulting maps portrayed the expected relative contributions of all marks at a given distance to an arbitrary predicted point (**Methods**).

**Figure 3.**
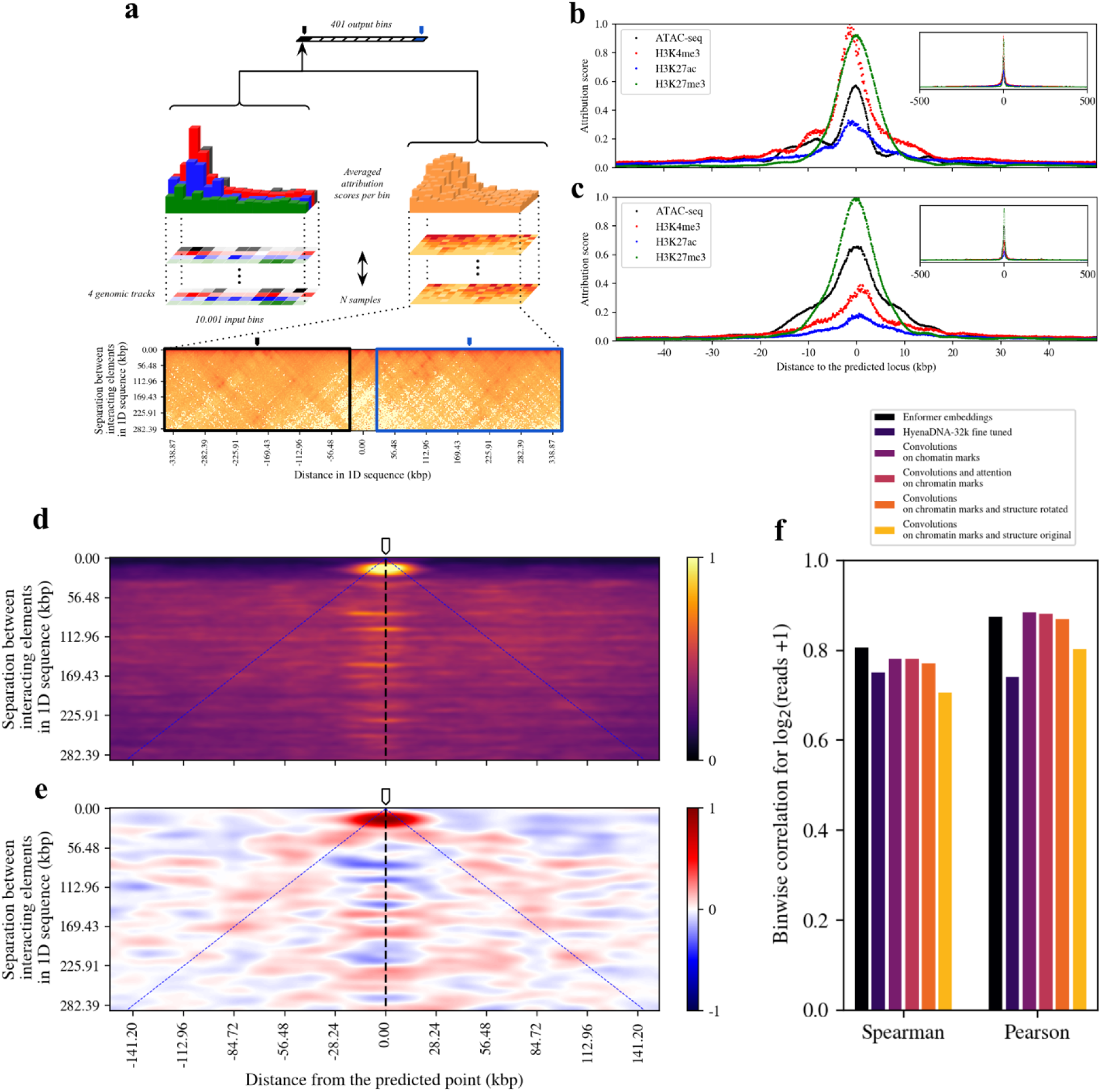
Averaged attributions provide information about chromatin mark significance and scale of influence in the prediction task. **a)** Attribution method schema. Micro-C arrays are rotated 45 degrees and corners are cropped to yield rectangular maps **(Supplementary Figure 8)**. Each of the positions in both chromatin arrays and structural arrays are given an attribution score towards the prediction of each single output node. Attribution arrays are centered, stacked and averaged to obtain an attribution map as in d). **b)** Averaged attributions of each genomic track in the chromatin landscape branch. Attributions are centered at the prediction site and rescaled between zero and the maximum attribution score. **c)** Chromatin landscape attributions when the model is also provided contact maps as input. **d)** Absolute attribution map for the chromatin structure branch. When overlaid to a contact map region like the blue or black rectangles in a), it conveys the importance that the model attributes on average to each part of the map when predicting nascent transcription levels at the genomic locus corresponding to the top center of the map (the pointer). Color scale ranges from no attribution, i.e. Score=0, to maximum attribution, with score 1. Map smoothed with gaussian kernel of σ=1. **e)** Average of signed attribution score maps for the structure branch. Scale ranges from -1 to 1, where 0 corresponds to no contribution. Positive values (red) are assigned to regions that contribute to raise output EU-seq levels. Negative values (blue) indicate an inverse relationship between input contact signals and output EU-seq levels. Map smoothed with a gaussian kernel of σ=2 to ease the identification of regions. **f)** Model performance comparison. The addition of contact information or attention to handle long range interactions did not improve the accuracy of the predictions.

The promoter mark H3K4me3 was found to be the most important predictor of nascent expression levels when using only the chromatin landscape branch of the model, as it was assigned the highest attribution score (i.e. feature importance) around the predicted point (**Figure 3b**). The averaged attribution scores of the silencing mark H3K27me3 were comparable to those of H3K4me3, while accessibility and the enhancer and promoter mark H3K27ac were assigned lower scores, peaking at around 60% and 40% of that of H3K4me3, respectively. Strikingly, the integrated gradients analysis presented a sharp, power-law decay of the attributions as a function of the distance to the predicted locus. Attribution scores dropped to less than 20% of their peak value after 5 to 10 kbp from the predicted locus, depending on the mark (**Figure 3b**). Here, the curves presented an elbow, after which the decay followed at a slower rate. Further than 50kbp away, the attribution scores were almost negligible. Thus, CLASTER relied mainly on a window of 20 kbp around the target locus to perform the predictions, while leaving most of the sequence unattended. This rather local behavior, observed also in the perturbational scenario, contrasts with the fact that 1Mbp of context was available to the network and provided a complementary validation to the results presented in [9]. Of note, a significantly wider field of view than 20kbp was covered by the dilated convolutions rather early in the network (**Supplementary Table 1)**, and therefore it is improbable that this locality appeared as a limitation of the receptive field of the convolutions.

### Micro-C maps provided an added layer of information and interpretability to CLASTER

The fact that the model was not attending to potential distal interactions motivated us to add high-resolution chromatin contact maps explicitly as inputs to the model. Convolutional neural networks could in principle be sensitive to various structural patterns in such contact matrices, e.g. point-like interactions between separated elements, chromatin compartments and topologically associated domains (TADs), which would not be explicitly accounted for in more classical approaches [10].

The use of contact maps in their usual, symmetrical format, induced in fact the creation of several convolutional filters capturing the aforementioned features (**Supplementary Figure 10, left)**. We found that the symmetry of the original matrices limited the power of image processing-based approaches when studying Micro-C data, as the nuanced features extracted in early stages of the network were lost in deeper layers, where primarily a broad sense of regional contact was propagated forward (**Supplementary Figure 10, right)**. In addition, complementing the available information with contact maps induced a decrease in the attributions of the model to the promoter mark (**Figure 3c**), suggesting a certain degree of overlap in the information coming from the two data modalities. To introduce contact information in a more appropriate manner, Micro-C matrices were rotated and cropped, and explored by scrolling the convolutional filters on the sequence axis only (**Figure 3a, Supplementary Figure 11, Methods**). This procedure reduced the redundancy of the information and skipped the contact-poor corner areas of the contact maps.

Following the same steps as before, we proceeded to evaluate the contributions of every bin in a Micro-C matrix towards the prediction of EU-seq levels at the central position (**Figure 3a**). The attribution map obtained from absolute scores brought to light two different behaviors of the network depending on the distance in 1D sequence between points in physical contact (**Figure 3d)**. Attributions of bins connecting loci closer than ca. 30 kbp showed a clear peak around the predicted point, i.e., the main contribution to the nascent transcription levels in the predicted locus came from the inputs describing the local environment and interactions of the predicted locus with its close neighborhood at a TAD scale. On the contrary, the network did not use the local neighborhood of loci located more than 30 kbp upstream or downstream of the prediction point. These results were in line with our previous findings based on chromatin state attributions. Interestingly, the network did now attend to regions of the map linking more distal regions. The network’s attributions for all bins connecting loci found at the same distance in 1D sequence were quite similar from 30 kbp on, yielding a certain positional invariance (**Figure 3d**). The main variation in attributions therefore occurred across different distances and was characterized by a sharp increase at around 30kbp.

To assess whether a high level of contact in a certain region of the map contributed to raising or lowering the expression levels at the predicted locus, we then computed averages of the maps with signed attributions (**Figure 3e, Supplementary Figures 12-13**). The neighborhood of the predicted point had once more the highest impact on the output, with a higher number of counts or contact in that region contributing the most to raising expression levels. Of note, positive attribution scores were assigned to loci in physical contact with the predicted locus, indicating also a positive association between input and output signals. This feature was quite conserved across contact distances from 30kbp onwards and suggested a potential boost of expression in the target profiles when a contact in such a region was to be found (**Figure 3e, blue lines)**. These results indicated that CLASTER could integrate contact information to that already provided in the chromatin landscape, understanding the impact of features in different regions of the contact maps towards the corresponding expression levels. The network now attended to distal interacting elements, but the main contributor to the predictions was still the local environment around the target locus, which displayed a clearly distinct behavior.

### Explicit modeling of distal interactions did not significantly impact the performance

The introduction of multi-headed attention has been shown to improve the prediction performance of DNA-based models [2,13,16]. Unlike convolutions, which process information at different scales hierarchically, the attention mechanism allows the models to process all parts of the sequence simultaneously. Chromatin-based models could also benefit from such a mechanism if the sequential appearance of patterns in the input or the interactions between distal regions were key factors to understand the data. We therefore introduced a Transformer Encoder block with two multiheaded attention layers, aiming to process the sequential appearance of the most abstract features in the chromatin landscape only setting (**Figure 1a, Supplementary Figure 1, Supplementary Table 1)**. However, a baseline model employing only convolutions on the chromatin landscape branch alone reached r_s_=0.7814 and r_p_=0.8852, performing as well as the one incorporating attention layers, which achieved r_s_=0.7815 and r_p_=0.8808. (**Figure 3f**). This was still the case when coupling high resolution structural information from Micro-C maps to CLASTER, with r_s_=0.7717 and r_p_=0.8690. Despite the model capturing rich features from structural data and understanding their relation to the output as shown previously, the added contact information did not boost the overall ability of CLASTER to fit the output EU-seq profiles.

Overall, our findings show that predictions could mostly be performed without relying on interactions between highly distal regulatory elements. The fact that dilated convolutions on chromatin states provided most of the required information to predict EU-seq profiles suggests that the chromatin landscape to RNA mapping problem might be better framed as a pattern recognition and matching task than a sequence processing one when predicting expression at a coarse, kilobasepair resolution. Finally, the final dense layers of CLASTER could already handle long-range interactions of abstract features, and hence the attention mechanism might need to be introduced earlier in the architecture to process lower-level features.

## Discussion

Our results support a rather confined gene regulation scenario. A predominantly local regulatory paradigm would seem plausible from the premises that most enhancers can activate most promoters [6,7] and that both frequency and intensity of the interactions between different genomic elements rapidly decay as a function of their separation in 1D sequence [17] and are further constrained by the 3D compartmentalization of chromatin. An increased body of work from different labs would in fact point in this direction from complementary perspectives. First, recent CRISPR-based studies have found variants in noncoding regions to alter mostly their nearmost genes [11,34–36] . Although predictive models have achieved high precisions in their predictions, the naïve baseline model associating a given genomic variant to the closest gene still provides a high sensitivity, i.e. a remarkably high number of true cases are captured under this assumption [34,37]. Global enhancer ablation assays in a native chromatin environment illustrated biases in the distribution of enhancers in the genome, showing a preference for cell type specific genes to be frequently located close to strong enhancers [6]. Finally, recent high resolution chromatin conformation capture assays proposed microcompartmentalization rather than loop extrusion mechanisms to be driving enhancer-promoter interactions [18], linking diseased phenotypes to the disruption of compartment boundaries [38].

Despite these findings, the difficulty of state-of-the-art machine learning approaches to capture the impact of distal enhancer variants in gene expression is still the focus of improvement for the next generation of models, which are rapidly benefiting from the latest advances in NLP and have a strong focus on the exploration of larger architectures [12] and the processing of longer context lengths [13,15,19,20]. These efforts contrast with the fact that early transformer-based models like the Enformer [2] already achieved remarkable results while relying on reduced genomic windows for the predictions [9]. Such behavior, which both the Enformer and CLASTER display, emerges as a consequence of the overall data distribution characterizing the studied system [6]. In other words, the spatial arrangement and organization of different regulatory elements in the genome does not reward the models enough to capture the expression changes induced by highly distal functional interactions, which become weaker [39] and increasingly rare with distance [9,40]. Despite long range interactions being reported to play crucial roles in specific regulatory processes [41–43], the results presented here support the idea that the information required to perform gene expression prediction is in most cases to be found in the close neighborhood of the predicted locus. Neither the addition of contact information matrices nor the introduction of the multi headed attention mechanism, both aimed to ease the processing of long-range interactions, yielded significant changes in overall performance. These findings, which build on top of previous work [6], have wide implications in the way we understand genomic organization and the extent to which it influences the transcriptional landscape, and invite us to rethink how new computational models can be designed to more suitably exploit such intrinsic properties of the genome.

We show how *in silico* perturbations of a given chromatin landscape can provide an added layer of explainability to the model’s predictions. These perturbative approaches exploit the rules in the input that the models have learned to explore the effects of unseen epigenetic variability but come with a number of challenges. On the one hand, perturbations of isolated input tracks can fail to mimic the expected biological behavior given the correlations established between the different marks, which define the chromatin states [29]. Chromatin-mark based models aimed to perform new synthetic/in-silico perturbations should either aim to replicate a complete desired chromatin state in a given locus of interest or successfully disentangle the roles of different marks. Sequence-based models might be less prone to display such correlational effects given that the one-hot encoding of nucleotides makes *in-silico* SNPs, insertions and deletions in-distribution alternative inputs for a given locus. However, the same type of correlational dynamics and co-appearance patterns might be learned between different parts of the sequence. In fact, Enformer has been shown to incorrectly predict the direction of the effects of certain genomic variants, in particular for non-monogenic traits that would imply the combined action of several loci [44,45]. These dynamics could constitute a hint towards the observation of a similar correlational learning framework as the one we reported in our first perturbational simulations, in this case across the sequence.

Left to be explored are better ways to couple high resolution contact matrices with chromatin mark profiles, with graph attention networks already showing great promise [19]. Nevertheless, the high sequencing depth requirement on higher resolution chromatin conformation maps and the related costs compromise scalability for machine-learning tasks. Our findings would motivate the exploration of region capture techniques to map shorter regions at a more fine-grained level [18]. Further work should aim to move from a mostly correlational learning framework to a more causal one, which is of utmost importance to determine the directionality of the association between transcription and the presence of certain chromatin marks. Training chromatin-based models in a GPT [46] or BERT [47] inspired masking fashion would perhaps provide the opportunity to do so, disentangling the roles of different epigenetic marks while studying sequences from varied cell type backgrounds. Finally, transcription factors and their binding profiles are a key missing piece of our models, with their addition as extra input tracks being the next logical step.

## Conclusions

We present CLASTER, a deep convolutional neural network that can be used to build an implicit gene regulation model, agnostic to DNA sequence and without relying on the manual extraction of a number of curated features. CLASTER is capable of understanding the relations established between different epigenetic marks describing a given chromatin state, how these states are combined to create a chromatin landscape, and how a given landscape and its matching three-dimensional arrangement can be translated into nascent transcription levels. Given its implementation using the EIR framework, CLASTER can easily be extended to new datasets, data modalities and tasks. Therefore, we expect our work to serve as an inspiration and an easy entry-point to epigenomics modeling for a broad range of researchers with diverse computational resources and scientific backgrounds.

## Methods

### Processing of chromatin mark array data

Normalized enrichment levels for ATAC-seq, H3K4me3, H3K27ac and H3K27me3 in mESCs were obtained at a 100bp resolution from publicly available data in the NCBI GEO under accession numbers **GSE146328** and **GSE186349**. Each sample consisted of an array of shape (N_chrom marks_, N_bins_) = (4, 10001) where each row was a different mark and each column a 100bp bin with the mark’s averaged enrichment level. Samples were centered at the TSS of a gene of interest. Note that a given gene could appear in different samples but at different locations, if it was close enough to the central predicted gene. A detailed description of the genomic regions constituting our samples and matching targets can be found in **Supplementary Table 2,** and a guide through data obtention and preprocessing steps can be found at the Github repository of the project https://github.com/RasmussenLab/CLASTER.

### Processing of nascent RNA array data

Publicly available nascent transcription profiles matching the input samples were obtained from **GSE146328** [27]. Since our aim was to estimate integrated nascent gene expression levels and to study the propagation and impact of *in silico* perturbations in as many genes as possible per sample, we prioritized the prediction of a wide genomic region over output resolution. 1Mbp regions were originally obtained at a resolution of 20 bp. A gaussian kernel of sigma = 50 was then applied to smooth the signals before averaging them in bins of 1 kbp (**Supplementary Figure 2**). This procedure reduced drastically the number of targets while preserving the main source of variation at the kbp scale, speeding up the computations of input-output attribution scores. The resulting profiles were symmetrically cropped to allow all predicted genes to have access to a wide chromatin range. Each target sample was finally chosen to be a region of 401 kbp centered at the TSS of the gene of interest. Target profiles were further cropped to fit the target windows of the Enformer, Hyenadna-160k and Hyenadna-32k, detailed below.

### Processing of micro-C contact maps

Micro-C matrices describing chromatin contact frequencies in mESCs were obtained at a resolution of 1600bp per bin from NCBI GEO under accession number **GSE130275** [28]. Lower resolution mcool representations started to lose the nuances in structural detail provided by MicroC, but higher resolution matrices became too sparse. Bin values had a range of 7 orders of magnitude after balancing, with a minimum of 6,807 · 10^-7^ and a maximum of 2,001 in our data. NaN values due to balancing and 0 count bins were then set to 1·10^-14^ → log_10_=-14. This arbitrarily small value was chosen to be the same distance apart from the minimum actual contact value as the minimum to maximum contact range. Imputing by the minimum 10^-7^ could be understood as an averaged homogeneous background contact, which we tried to avoid by giving those bins a clearly distinct behavior. Micro-C matrices were originally treated as 2D arrays of shape (625, 625) containing log_10_ transformed bin counts. A list of regions for which Micro-C matching maps could not be obtained is found in **Supplementary Table 5**.

### Rotations and cropping of Micro-C maps

Contact matrices displayed two properties: 1) they were symmetrical and 2) the number of reads decreased when exploring regions far from the diagonal. Treating them as images, i.e. allowing for the convolutions to scroll in both x and y directions, led to an under usage of the model’s capabilities given the redundancy found in both triangles and the lack of information provided by the regions close to the corners far from the diagonal. In order to provide the contact information in a more appropriate manner for a model such as CLASTER, Micro-C matrices were rotated 45 degrees and cropped. This procedure allowed us to 1) reduce the number of inputs, accelerating the computations and 2) define deeper 1D kernels exploring all distances at once. These kernels were then only allowed to scroll sideways, in the original diagonal direction. Given the symmetry of the matrices, we kept only the upper triangle. The rotated triangle was cropped to ca. 0.207 times the original length to create a rectangular matrix of shape (129, 626). Distances and scales in the new matrix are illustrated in **Supplementary Figure 11** and can be expressed as follows: Expressing a and a’ in terms of L, we have:

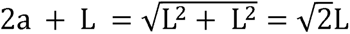

We get 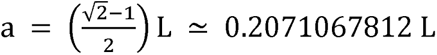 and 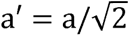. After rotating and cropping the matrices, we have 626 bins for a smaller window of observation. The width of the new window was then 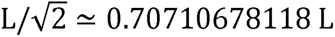 where L - 1 Mbp. Linear bin densities in the original and transformed Micro-C matrices were:

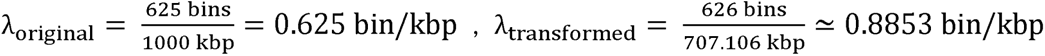

The interaction distance d_1D_, i.e. the real separation in 1D sequence between two interacting elements, was implicitly encoded in the new y coordinate y_new_, which was the distance in the perpendicular direction to the diagonal. A point at a distance y_new_ from the diagonal, found at a column bin y*_bins_* in the rotated and cropped matrices, connected the points in the diagonal at -x_bins_ = -|y_bins_| and x_bins_ = |y_bins_| given that the scale was the same in both axes. This distance can be converted from new bins to kbp in 1D sequence using the diagonal bin density found before as 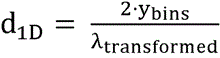.

### Reference genome, gene annotations and enhancer annotations

The mouse reference genome and protein coding gene annotations were downloaded from the GENCODE (mouse; GRCm38 release 25) [48]. Proximal and distal Enhancer-Like Signatures (pELS, dELS) in mice were obtained from **Supplementary Table 11** in [30].

### In silico perturbations

In the first perturbational scenario (P1), the loss of enhancer activity was simulated by setting to zero reads the levels of the enhancer mark H3K27ac, for all enhancers i - 1,…,N_enh_ inside the target window, enhancer by enhancer. In the second perturbational scenario (P2), a heterochromatin state substitution was performed for all bins inside the enhancer boundaries by setting the levels of all marks to 0 reads except from H3K27me3. This arbitrary silenced-like state was aimed to remove the contributions of accessibility, background promoter and enhancer activities while keeping a plausible level of H3K27me3 in the corresponding chromatin environment. We then measured the relative change in averaged nascent transcription that the reference gene in each sample experienced when ablating every single active enhancer falling inside the predicted region of the genome. The relative change between baseline and perturbed input predictions was computed as:

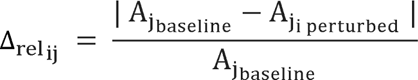

Here, A_jbaseline_ and A_ji perturbed_ described the area under the EU-seq curves inside the boundaries of reference gene j normalized by that gene’s length, when predicting the output with the control chromatin landscape and the output from the same chromatin landscape with the i_th_ enhancer ablated. To integrate the area under reference genes shorter than 1 kbp or spanning more than the prediction window’s boundaries we used widths of 1 bin and half of the prediction window’s length, since the TSS was located in the center. Only active enhancers were kept for the analyses, where an active enhancer was defined as a pELS or dELS region with an average enrichment of H3K27ac above a predefined threshold, set to be 10 reads. Enhancer gene pairs for reference genes with zero reads were discarded from the analyses. Since the background distribution of number of genes between enhancers and target genes was not uniform, histograms of the most significant enhancer-gene pairs per sample were shown unnormalized (in the main manuscript) and normalized by the background distribution of pairs present in the test dataset (in the supplementary, **Supplementary Figure 9**).

### Scoring the contributions of each relative input location to the output

To score the contributions of a given input position towards the expression levels in a target locus, we used the widespread integrated gradients axiomatic attribution method [33] as implemented in the Captum python library. Integrated gradients were computed as the path integral of the gradients along the straight line path connecting a predefined baseline signal x’ and a given input x. The baseline background signal was obtained by averaging 256 samples in each modality. The authors define the integrated gradient along the i_th_ dimension as:

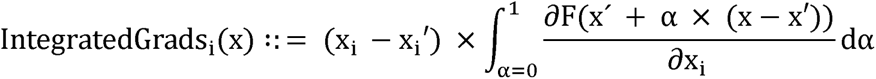

As stated before, the chromatin landscape inputs were described by a matrix of shape (N_chrom marks_, N_input bins_) = (4, 10001) exploring 1Mbp in bins of 100bp. Chromatin structure was given by a matrix of shape (129, 626) describing the same region at a resolution of 1.6 kbp. EU-seq targets were described by a vector of length N_output bins_ = 401 exploring from -200kbp to 200kbp relative to the TSS of a gene of interest in bins of 1 kbp. Signed and absolute attribution scores were then obtained for every bin position in the input arrays towards the prediction of every target position (**Supplementary Figure 12**). To obtain averages over predicted points, attribution maps for each target node were then rolled in steps of 10 bins = 1kbp to locate the predicted target to the center of the map. High frequency noise arising from the translations of the maps was removed by adding a small uncertainty in the number of bins to roll for the chromatin landscape, adding eps number of bins with -10 bins < eps < +10 bins. A similar smoothing effect was achieved for structural attributions by applying a gaussian filter (**Supplementary Figure 13**). The maps were then symmetrically cropped to keep only the relative positions to the predicted point that had been measured in all predictions, with measurements centered at -200 kbp and 200 kbp setting the limits (**Supplementary Figure 9**).

### CLASTER

The architecture of CLASTER is illustrated in **Supplementary Figure 1** and further detailed in **Supplementary Table 1**. In order to translate the input to the corresponding output, features were extracted separately from both branches of the model, namely the chromatin landscape branch and the chromatin structure branch but using the same principles. These include a set of layers implementing dilated convolutions, residual connections and optional attention heads. Input dimensionality reduction was achieved by scrolling the convolutional kernels in strides of 2. The extracted features at different scales were integrated at a later stage in the network, the fusion module. Further dimensionality reduction and mapping to the output nascent RNA levels was performed with a final set of densely connected layers (MLP). The MLP ended in a hidden vector representation, and finally each node was linearly connected to every single prediction point, i.e 401 bins at 1kbp resolution. A final non-trainable ReLU was applied to cut negative values. Configuration files to build and run the models using the EIR framework [26] in both training and test settings are available at https://github.com/RasmussenLab/CLASTER.

### Training

The network was trained in batches of 64 samples and a learning rate of 1e-4. We used the AdamW optimizer with default parameters [49]. Regularization measures included the use of high dropout values in the MLP layers, setting a given stochastic depth (probability to remove residual block) and the use of SmoothL1 as the loss function. Flipped samples were also added to the training set for data augmentation purposes but were not used for the test set analyses. Chr17 samples were used as a hold out validation, and Chr4 was used as a hold out test. A detailed description of the genes of interest that were used to create our samples can be found in **Supplementary Table 2**. Samples for which we could not obtain Micro-C matching maps were removed from the training, validation or test sets in all settings, and can be found in **Supplementary Table 4**.

### Benchmarks

Sequence embeddings corresponding to the central 160 kbp of the genomic regions of study were obtained from the backbone of HyenaDNA-160k [15] with pretrained, frozen weights. Code to build HyenaDNA-160k was obtained from the publicly accessible google colab [50] and pretrained weights were obtained from [51]. The input sequences when using HyenaDNA-160kbp ranged in this case from -80kbp to 80kbp after the TSS of the central gene in the observation window. We then trained a simple model or head on top of the frozen embeddings to predict our EU-seq signals at the same resolution as we did with CLASTER (**Supplementary Figures 3-4)**. The head performed an average pooling over the sequence axis to reduce sequence length at a 128 bp resolution. To ease the predictions, a first set of convolutions of kernel 9 and dilation 2 were applied to the resulting embeddings. This enabled the head to identify patterns at a scale of ca. 2 kbp. These patterns were then mapped to the targets using a dense layer.

We tried to perform the regression of the target EU-Seq profiles using the pretrained embeddings from the HyenaDNA-160k, but we encountered a number of challenges. First, in the original manuscript the authors performed mainly classification tasks at a sequence level, but simultaneous regressions of multiple outputs were not explored. Second, HyenaDNA was pretrained on human and not mouse genomic sequences, although we would not expect this to be a core issue. We believe that the pretrained embeddings obtained by next token prediction did not encode enough functional information to perform the EU-Seq predictions, since further fine-tunning of the model was required for our downstream task. This was importantly not the case with Enformer-based embeddings. This fine-tuning requirement might not be unique to HyenaDNA. Finally, Hyena’s design was aimed to keep nucleotide resolution, but this would not be necessary to predict averages of read counts at a coarse, kbp resolution. Despite the low parameter count and high efficiency of the Hyena operator, which reduced drastically the number of trainable parameters, the network’s design yielded a high number of inner sequence representations, the activations at each filter in each layer, that kept sequence length (160,000 columns) while adding an embedding dimension (256 rows, e.g.). Each single sequence representation as a matrix of float32 values required:

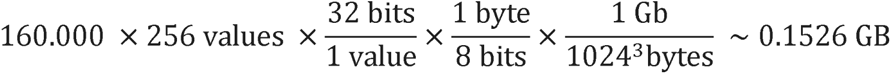

with many of them being stored already in the forward pass. This imposed a significant challenge, rapidly increasing memory and computing time requirements, which constrained us to fine-tune Hyena-160k in batches of only 2 sequences in an NVIDIA A100 GPU with 80GB of RAM. This motivated us to use the smaller Hyena-32k model, for which we could load the pretrained weights and fine-tune the pretrained backbone together with the added head in batches of 16 samples in a reasonable period of time. Targets were cropped to match the new context length to the range -15kbp to +15kbp.

The same procedure was repeated for the Enformer architecture [2]. Sequence embeddings corresponding to the central 160kbp of the genomic regions of study were obtained from the backbone of Enformer [2] with pretrained, frozen weights. Code to build the Enformer in Pytorch and matching pretrained weights were obtained from [52]. To match the previous scenario when using the Enformer, we used the same 160kbp sequences as in Hyena and symmetrically prepended and appended N nucleotides as proposed in [9], which acted as masks, to provide the same available information in both settings. The embeddings of the Enformer, of shape (1, 896, 3072) corresponded to 896 × 128 bp - 114688 bp ∼ 115 kbp. The matching targets were therefore cropped to keep the range between -57kbp to +57kbp from the central gene’s TSS to match the Enformer’s prediction window.

## Availability of data and materials

The raw data used in this project was obtained from publicly available sources [27,28,32]. The original bigwig files for chromatin mark enrichment, ATAC-seq and EU-seq can be found at NCBI’s GEO under accession numbers **GSE146328** and **GSE186349**. Mcool files storing high resolution chromatin contact frequency maps can be found under the accession number **GSE130275**. Processed files, input and target arrays and configuration files to build CLASTER and reproduce our analyses can be obtained following the python notebooks found at the project’s Github repository https://github.com/RasmussenLab/CLASTER.

All code is available at the github repository of the project. EIR can be found at https://github.com/arnor-sigurdsson/EIR. The code of the project was organized in a number of python notebooks covering 0) a tutorial, 1) how to download the raw data from NCBI and preprocess it to create inputs and targets for EIR-based models, 2) how to create, train and test CLASTER using EIR, 2b) how to add custom heads on top of Enformer or HyenaDNA embeddings and fine-tune HyenaDNA-32k with an added custom regression head and finally 3) how to perform the data analysis and create the figures presented in this manuscript.

## Acknowledgments

We would like to thank Nils Krietenstein for his valuable input and feedback on the project. We would also like to acknowledge the members of the Rasmussen lab, in particular Justus Gräf and Henry Webel, for our fruitful discussions. We would finally like to thank the Novo Nordisk Foundation Center for Protein Research, the Novo Nordisk Foundation Center for Basic Metabolic Research and the Novo Nordisk Foundation Center for Genomic Mechanisms of Disease for their support.

## Competing interests

S.R. is the founder and owner of BioAI. The remaining authors declare no conflicts of interest.

## Funding

M.P.A was funded by the Novo Nordisk Foundation Copenhagen Bioscience Ph.D. Program grant No. NNF22SA0078231. M.P.A, A.S., T.N., N.K, C.C, and S.R. were supported by the Novo Nordisk Foundation (NNF14CC0001). M.P.A., A.S. and S.R. were supported by the Novo Nordisk Foundation (NNF23SA0084103 and NNF21SA0072102). C.C. was supported by the Novo Nordisk Foundation (nos. NNF20OC0065482 and NNF22OC0074677). The Novo Nordisk Foundation Center for Protein Research is financially supported by the Novo Nordisk Foundation (no. NNF14CC0001).

## Author Contributions

S.R designed and supervised the project and wrote the manuscript. C.C designed and supervised the project. M.P.A wrote the code, carried out the analyses, interpreted the results, and wrote the manuscript. A.S wrote the code and assisted in the analyses. T.N acquired and preprocessed the data and assisted in the analyses. All authors reviewed and edited the final manuscript.

## Supplementary information

**Additional file 1.** Supplementary Figures 1 to 13.

**Additional file 2.** Supplementary Tables 1 to 5.

## Notes

### Competing Interest Statement

Simon Rasmussen is the owner and cofounder of a Biotech AI consulting company (BioAI).

## References

1. Zhou J, Theesfeld CL, Yao K, Chen KM, Wong AK, Troyanskaya OG. Deep learning sequence-based ab initio prediction of variant effects on expression and disease risk. Nat Genet. 2018;50:1171–9.

2. Avsec Ž, Agarwal V, Visentin D, Ledsam JR, Grabska-Barwinska A, Taylor KR, et al. Effective gene expression prediction from sequence by integrating long-range interactions. Nat Methods. 2021;18:1196–203.

3. Avsec Ž, Weilert M, Shrikumar A, Krueger S, Alexandari A, Dalal K, et al. Base-resolution models of transcription-factor binding reveal soft motif syntax. Nat Genet. 2021;53:354–66.

4. Kelley DR, Reshef YA, Bileschi M, Belanger D, McLean CY, Snoek J. Sequential regulatory activity prediction across chromosomes with convolutional neural networks. Genome Res. 2018;28:739–50.

5. Chen KM, Wong AK, Troyanskaya OG, Zhou J. A sequence-based global map of regulatory activity for deciphering human genetics. Nat Genet. 2022;54:940–9.

6. Narita T, Higashijima Y, Kilic S, Maskey E, Neumann K, Choudhary C. The logic of native enhancer-promoter compatibility and cell-type-specific gene expression variation [Internet]. bioRxiv. 2022 [cited 2023 Sep 1]. p. 2022.07.18.500456. Available from: https://www.biorxiv.org/content/biorxiv/early/2022/07/19/2022.07.18.500456

7. Bergman DT, Jones TR, Liu V, Ray J, Jagoda E, Siraj L, et al. Compatibility rules of human enhancer and promoter sequences. Nature. 2022;607:176–84.

8. Hong CKY, Cohen BA. Genomic environments scale the activities of diverse core promoters. Genome Res. 2022;32:85–96.

9. Karollus A, Mauermeier T, Gagneur J. Current sequence-based models capture gene expression determinants in promoters but mostly ignore distal enhancers. Genome Biol. 2023;24:56.

10. Fulco CP, Nasser J, Jones TR, Munson G, Bergman DT, Subramanian V, et al. Activity-by-contact model of enhancer–promoter regulation from thousands of CRISPR perturbations. Nat Genet. 2019;51:1664–9.

11. Gschwind AR, Mualim KS, Karbalayghareh A, Sheth MU, Dey KK, Jagoda E, et al. An encyclopedia of enhancer-gene regulatory interactions in the human genome [Internet]. bioRxiv. 2023 [cited 2023 Nov 21]. p. 2023.11.09.563812. Available from: https://www.biorxiv.org/content/biorxiv/early/2023/11/13/2023.11.09.563812

12. Dalla-Torre H, Gonzalez L, Mendoza-Revilla J, Carranza NL, Grzywaczewski AH, Oteri F, et al. The Nucleotide Transformer: Building and Evaluating Robust Foundation Models for Human Genomics [Internet]. bioRxiv. 2023 [cited 2023 Nov 2]. p. 2023.01.11.523679. Available from: https://www.biorxiv.org/content/biorxiv/early/2023/09/19/2023.01.11.523679

13. Linder J, Srivastava D, Yuan H, Agarwal V, Kelley DR. Predicting RNA-seq coverage from DNA sequence as a unifying model of gene regulation [Internet]. bioRxiv. 2023 [cited 2023 Nov 2]. p. 2023.08.30.555582. Available from: https://www.biorxiv.org/content/biorxiv/early/2023/09/01/2023.08.30.555582

14. Ji Y, Zhou Z, Liu H, Davuluri RV. DNABERT: pre-trained Bidirectional Encoder Representations from Transformers model for DNA-language in genome. Bioinformatics. 2021;37:2112–20.

15. Nguyen E, Poli M, Faizi M, Thomas A, Birch-Sykes C, Wornow M, et al. HyenaDNA: Long-Range Genomic Sequence Modeling at Single Nucleotide Resolution [Internet]. arXiv [cs.LG]. 2023. Available from: http://arxiv.org/abs/2306.15794

16. Nguyen E, Poli M, Durrant MG, Thomas AW, Kang B, Sullivan J, et al. Sequence modeling and design from molecular to genome scale with Evo [Internet]. bioRxiv. 2024 [cited 2024 Mar 11]. p. 2024.02.27.582234. Available from: https://www.biorxiv.org/content/10.1101/2024.02.27.582234v2

17. Krietenstein N, Abraham S, Venev SV, Abdennur N, Gibcus J, Hsieh T-HS, et al. Ultrastructural details of mammalian chromosome architecture. Mol Cell. 2020;78:554–65.e7.

18. Goel VY, Huseyin MK, Hansen AS. Region Capture Micro-C reveals coalescence of enhancers and promoters into nested microcompartments. Nat Genet. 2023;55:1048–56.

19. Karbalayghareh A, Sahin M, Leslie CS. Chromatin interaction-aware gene regulatory modeling with graph attention networks. Genome Res. 2022;32:930–44.

20. Bigness J, Loinaz X, Patel S, Larschan E, Singh R. Integrating Long-Range Regulatory Interactions to Predict Gene Expression Using Graph Convolutional Networks. J Comput Biol. 2022;29:409–24.

21. Singh R, Lanchantin J, Robins G, Qi Y. DeepChrome: deep-learning for predicting gene expression from histone modifications. Bioinformatics. 2016;32:i639–48.

22. Chen Y, Xie M, Wen J. Predicting gene expression from histone modifications with self-attention based neural networks and transfer learning. Front Genet. 2022;13:1081842.

23. Yang R, Das A, Gao VR, Karbalayghareh A, Noble WS, Bilmes JA, et al. Epiphany: predicting Hi-C contact maps from 1D epigenomic signals. Genome Biol. 2023;24:134.

24. Feng F, Yao Y, Wang XQD, Zhang X, Liu J. Connecting high-resolution 3D chromatin organization with epigenomics. Nat Commun. 2022;13:2054.

25. Cappelluti MA, Mollica Poeta V, Valsoni S, Quarato P, Merlin S, Merelli I, et al. Durable and efficient gene silencing in vivo by hit-and-run epigenome editing. Nature [Internet]. 2024; Available from: 10.1038/s41586-024-07087-8

26. Sigurdsson AI, Louloudis I, Banasik K, Westergaard D, Winther O, Lund O, et al. Deep integrative models for large-scale human genomics. Nucleic Acids Res. 2023;51:e67.

27. Narita T, Ito S, Higashijima Y, Chu WK, Neumann K, Walter J, et al. Enhancers are activated by p300/CBP activity-dependent PIC assembly, RNAPII recruitment, and pause release. Mol Cell. 2021;81:2166–82.e6.

28. Hsieh T-HS, Cattoglio C, Slobodyanyuk E, Hansen AS, Rando OJ, Tjian R, et al. Resolving the 3D Landscape of Transcription-Linked Mammalian Chromatin Folding. Mol Cell. 2020;78:539–53.e8.

29. Ernst J, Kellis M. Chromatin-state discovery and genome annotation with ChromHMM. Nat Protoc. 2017;12:2478–92.

30. ENCODE Project Consortium, Moore JE, Purcaro MJ, Pratt HE, Epstein CB, Shoresh N, et al. Expanded encyclopaedias of DNA elements in the human and mouse genomes. Nature. 2020;583:699–710.

31. Huang C, Helin K. Catching active enhancers via H2B N-terminal acetylation. Nat. Genet. 2023. p. 525–6.

32. Narita T, Higashijima Y, Kilic S, Liebner T, Walter J, Choudhary C. Acetylation of histone H2B marks active enhancers and predicts CBP/p300 target genes. Nat Genet. 2023;55:679–92.

33. Sundararajan M, Taly A, Yan Q. Axiomatic Attribution for Deep Networks [Internet]. arXiv [cs.LG]. 2017. Available from: http://arxiv.org/abs/1703.01365

34. Nasser J, Bergman DT, Fulco CP, Guckelberger P, Doughty BR, Patwardhan TA, et al. Genome-wide enhancer maps link risk variants to disease genes. Nature. 2021;593:238–43.

35. Morris JA, Caragine C, Daniloski Z, Domingo J, Barry T, Lu L, et al. Discovery of target genes and pathways at GWAS loci by pooled single-cell CRISPR screens. Science. 2023;380:eadh7699.

36. Gasperini M, Hill AJ, McFaline-Figueroa JL, Martin B, Kim S, Zhang MD, et al. A Genome-wide Framework for Mapping Gene Regulation via Cellular Genetic Screens. Cell. 2019;176:1516.

37. Mountjoy E, Schmidt EM, Carmona M, Schwartzentruber J, Peat G, Miranda A, et al. An open approach to systematically prioritize causal variants and genes at all published human GWAS trait-associated loci. Nat Genet. 2021;53:1527–33.

38. Kim KL, Rahme GJ, Goel VY, El Farran CA, Hansen AS, Bernstein BE. Dissection of a CTCF topological boundary uncovers principles of enhancer-oncogene regulation. Mol Cell [Internet]. 2024; Available from: 10.1016/j.molcel.2024.02.007

39. Zuin J, Roth G, Zhan Y, Cramard J, Redolfi J, Piskadlo E, et al. Nonlinear control of transcription through enhancer-promoter interactions. Nature. 2022;604:571–7.

40. Michaud EJ, Liu Z, Girit U, Tegmark M. The Quantization Model of Neural Scaling. Thirty-seventh Conference on Neural Information Processing Systems [Internet]. 2023. Available from: https://openreview.net/forum?id=3tbTw2ga8K

41. Lettice LA, Heaney SJH, Purdie LA, Li L, de Beer P, Oostra BA, et al. A long-range Shh enhancer regulates expression in the developing limb and fin and is associated with preaxial polydactyly. Hum Mol Genet. 2003;12:1725–35.

42. Uslu VV, Petretich M, Ruf S, Langenfeld K, Fonseca NA, Marioni JC, et al. Long-range enhancers regulating Myc expression are required for normal facial morphogenesis. Nat Genet. 2014;46:753–8.

43. Claussnitzer M, Dankel SN, Kim K-H, Quon G, Meuleman W, Haugen C, et al. FTO Obesity Variant Circuitry and Adipocyte Browning in Humans. N Engl J Med. 2015;373:895–907.

44. Sasse A, Ng B, Spiro AE, Tasaki S, Bennett DA, Gaiteri C, et al. Benchmarking of deep neural networks for predicting personal gene expression from DNA sequence highlights shortcomings. bioRxiv [Internet]. 2023; Available from: 10.1101/2023.03.16.532969

45. Huang C, Shuai RW, Baokar P, Chung R, Rastogi R, Kathail P, et al. Personal transcriptome variation is poorly explained by current genomic deep learning models. Nat Genet. 2023;55:2056–9.

46. Radford A, Wu J, Child R, Luan D, Amodei D, Sutskever I. Language Models are Unsupervised Multitask Learners. OpenAI [Internet]. 2019; Available from: https://dcmpx.remotevs.com/net/cloudfront/d4mucfpksywv/SL/better-language-models/language_models_are_unsupervised_multitask_learners.pdf

47. Devlin J, Chang M-W, Lee K, Toutanova K. BERT: Pre-training of Deep Bidirectional Transformers for Language Understanding [Internet]. arXiv [cs.CL]. 2018. Available from: http://arxiv.org/abs/1810.04805

48. Frankish A, Diekhans M, Jungreis I, Lagarde J, Loveland JE, Mudge JM, et al. GENCODE 2021. Nucleic Acids Res. 2021;49:D916–23.

49. Loshchilov I, Hutter F. Decoupled Weight Decay Regularization. International Conference on Learning Representations [Internet]. 2019. Available from: https://openreview.net/forum?id=Bkg6RiCqY7

50. Google colab [Internet]. [cited 2024 May 15]. Available from: https://colab.research.google.com/drive/1wyVEQd4R3HYLTUOXEEQmp_I8aNC_aLhL

51. LongSafari [Internet]. [cited 2024 May 15]. Available from: https://huggingface.co/LongSafari

52. Wang P. enformer-pytorch: Implementation of Enformer, Deepmind’s attention network for predicting gene expression, in Pytorch [Internet]. Github; [cited 2024 May 15]. Available from: https://github.com/lucidrains/enformer-pytorch

