## Supplementary Figures for "Modeling nascent transcription from chromatin landscape and structure"

**Affiliations**

**Author List Footnotes**

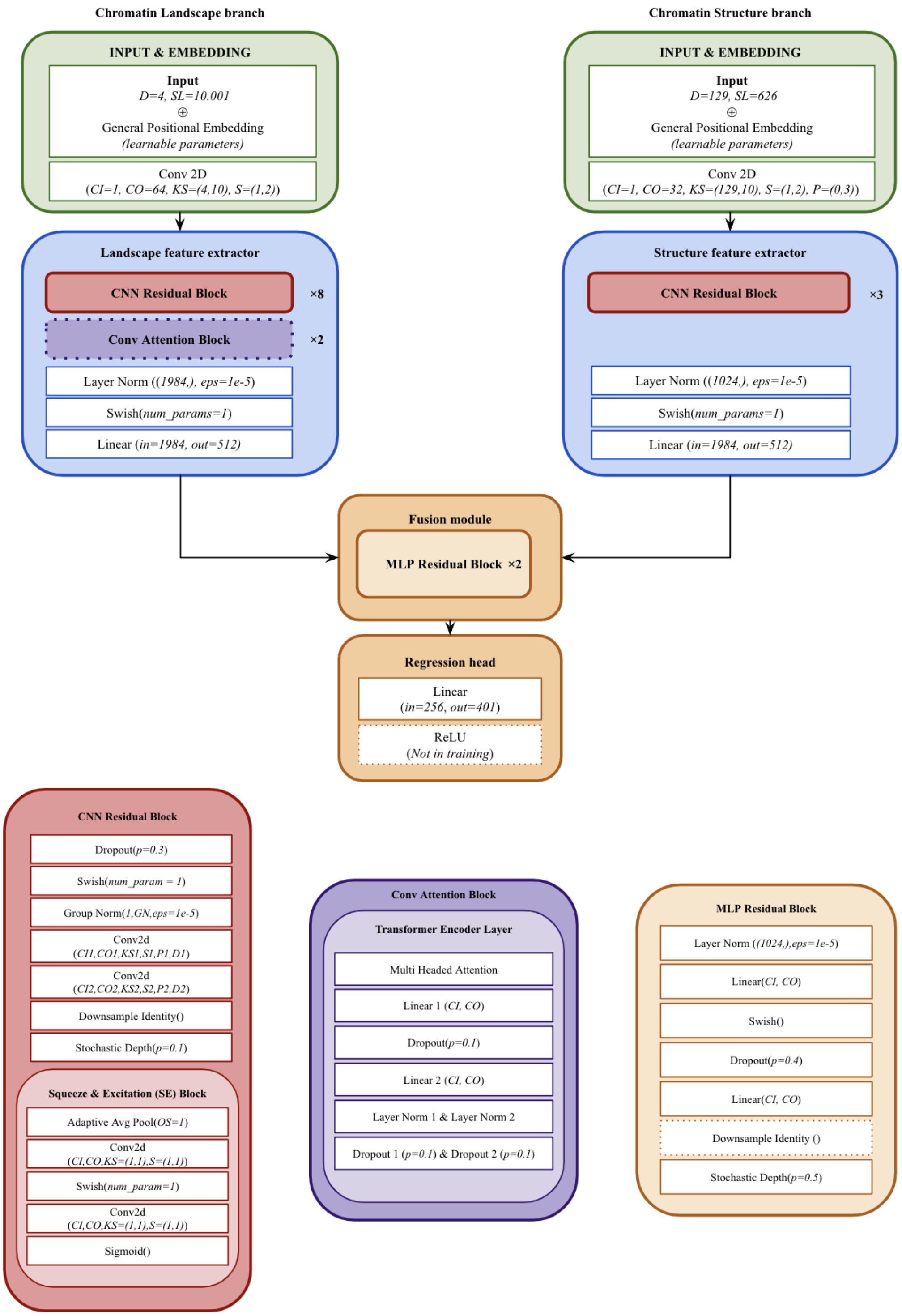


**Supplementary Figure 1. CLASTER architecture.** Specific values for Channels In (CI), Channels Out (CO), Kernel Size (KS), Stride (S), Padding (P), Dilation(D), etc. for specific layers and models are detailed in **Supplementary Table 1**. Configuration files to build the model using the EIR framework and a detailed description of each layer can be obtained from the Github repository of the project.

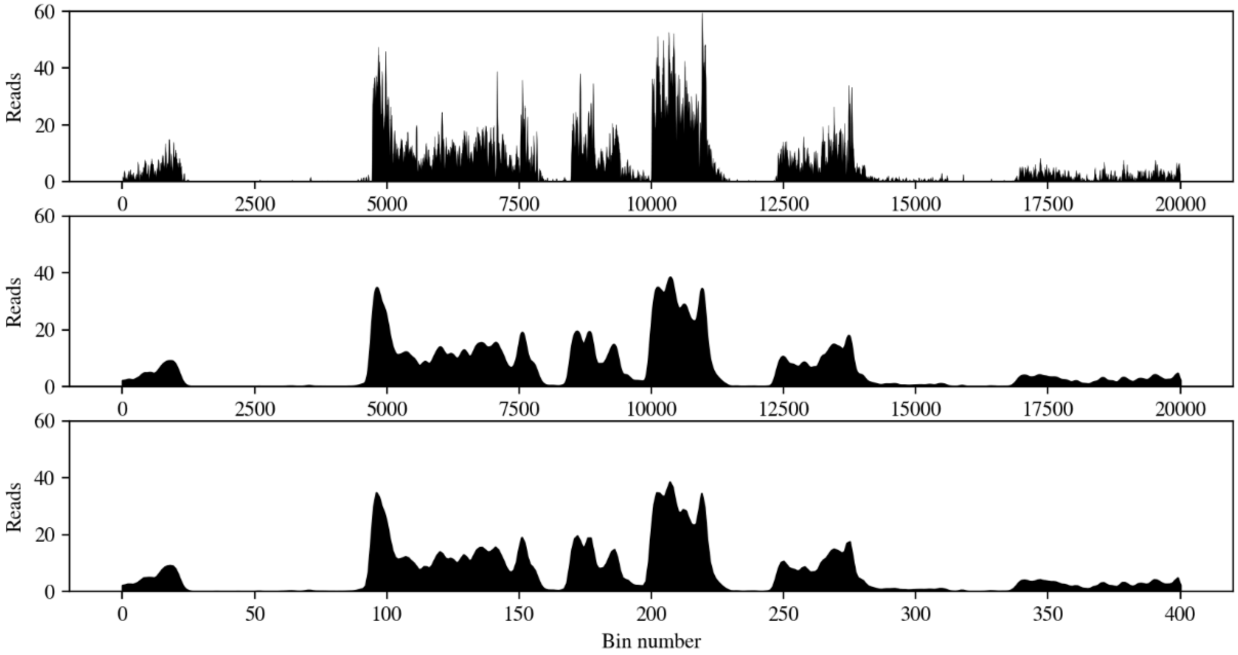


**Supplementary Figure 2**: **EU-seq profile smoothing and downsizing preserves the main source of variation at the kbp scale while reducing the number of targets. a)** Original EU-seq profiles at a 20bp resolution around the gene Nol9 (chr4). **b)** Smoothed profiles (σ=50 and bins of 20bp) keeping the original sampling rate. **c)** Final profile sampling in steps of 50 bins. Reducing the number of targets reduced drastically the number of input-target attribution scores to compute.

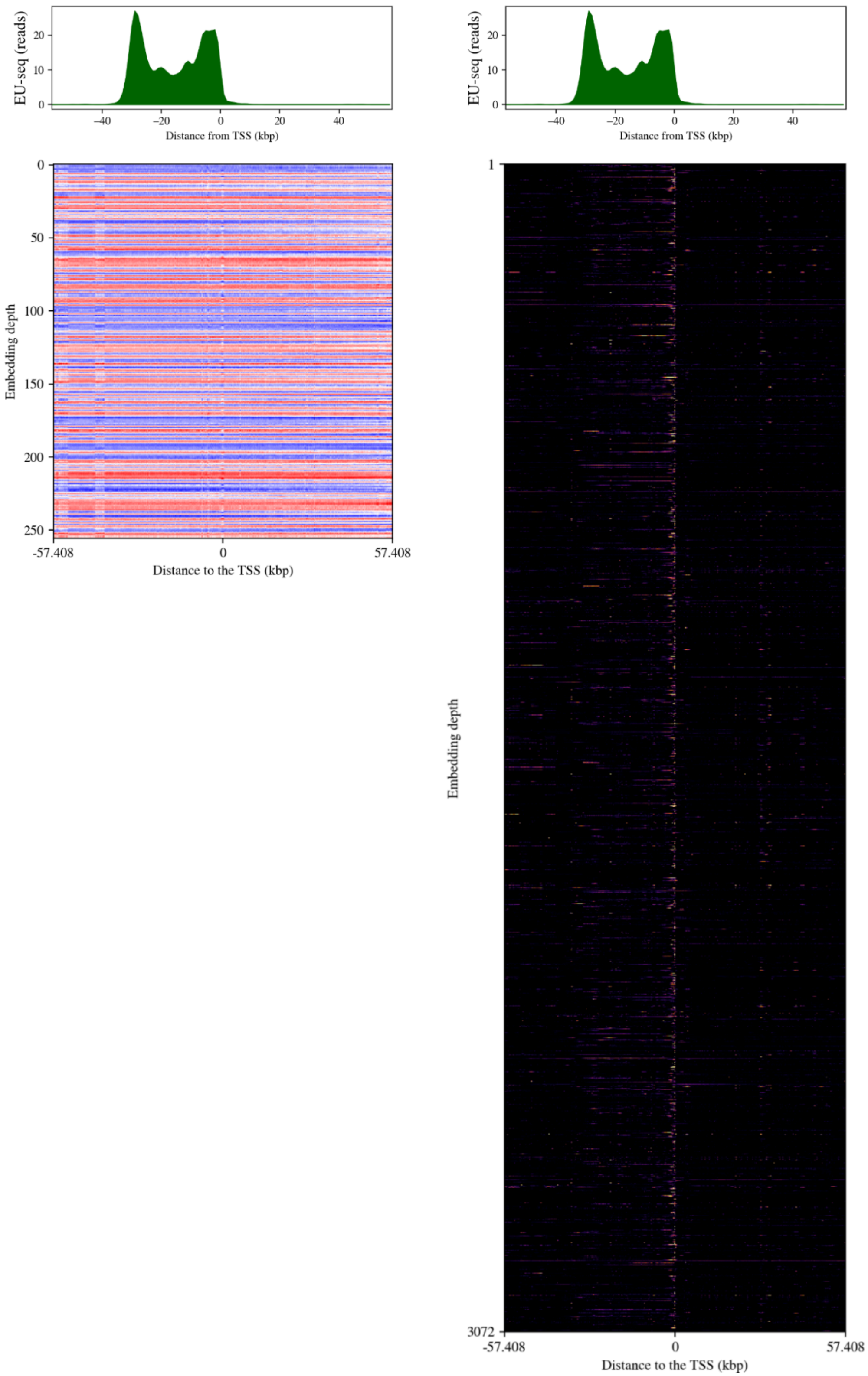


**Supplementary Figure 3. HyenaDNA-160k and Enformer pretrained embeddings encode varying degrees of functional information.** a) HyenaDNA-medium-160k-seqlen pretrained embeddings for the 160kbp around the gene Impad1 (ENSMUSG00000066324.2, chr 4). b) Matching embeddings from the pretrained Enformer backbone.

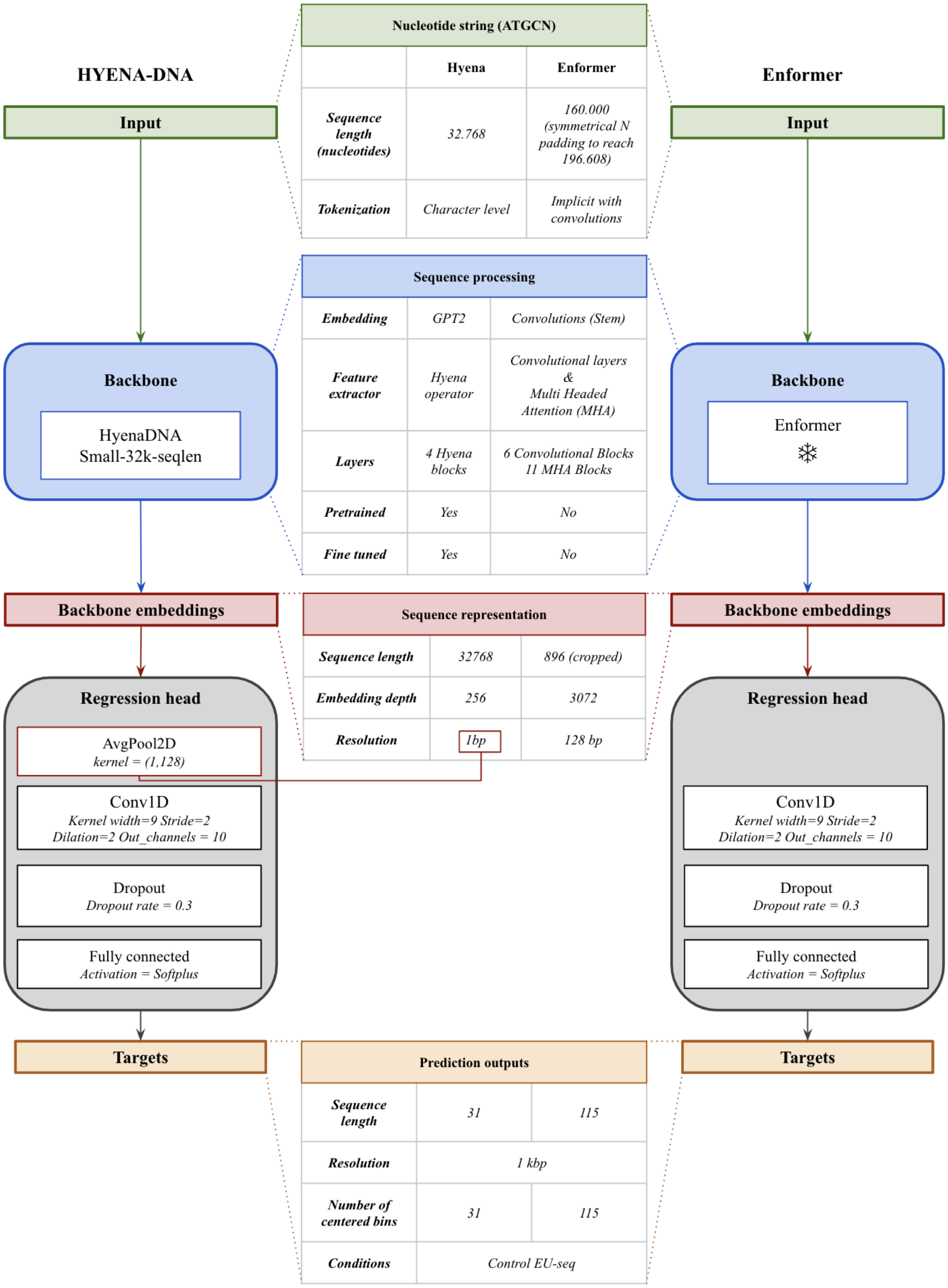


**Supplementary Figure 4. Enformer and HyenaDNA backbones & regression heads.** Note that an average pooling over the sequence axis was performed to Hyena's backbone embeddings to match the bin size of the Enformer. A kernel width of 9 with dilation of 2 provided enough context (ca. 2kbp) per convolution, which was further combined in the final Fully Connected layer to ease the predictions.

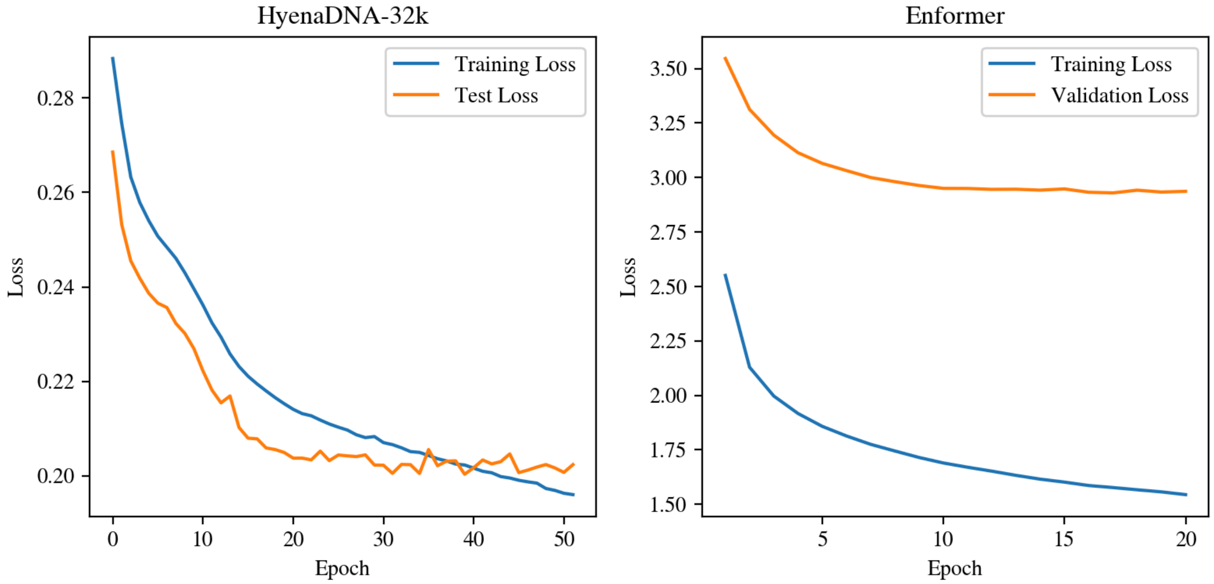


**Supplementary Figure 5. Loss curves for the benchmark model heads. Left)** HyenaDNA-32k loss curve when loading the pretrained backbone weights and fine tuning the combined model. **Right)** Loss curve when training a model head on top of the Enformer's pretrained embeddings. Both heads were identical except from a first average pooling (128bp) over the sequence axis for HyenaDNA-32k and a different number of target nodes to fit their respective context lengths.

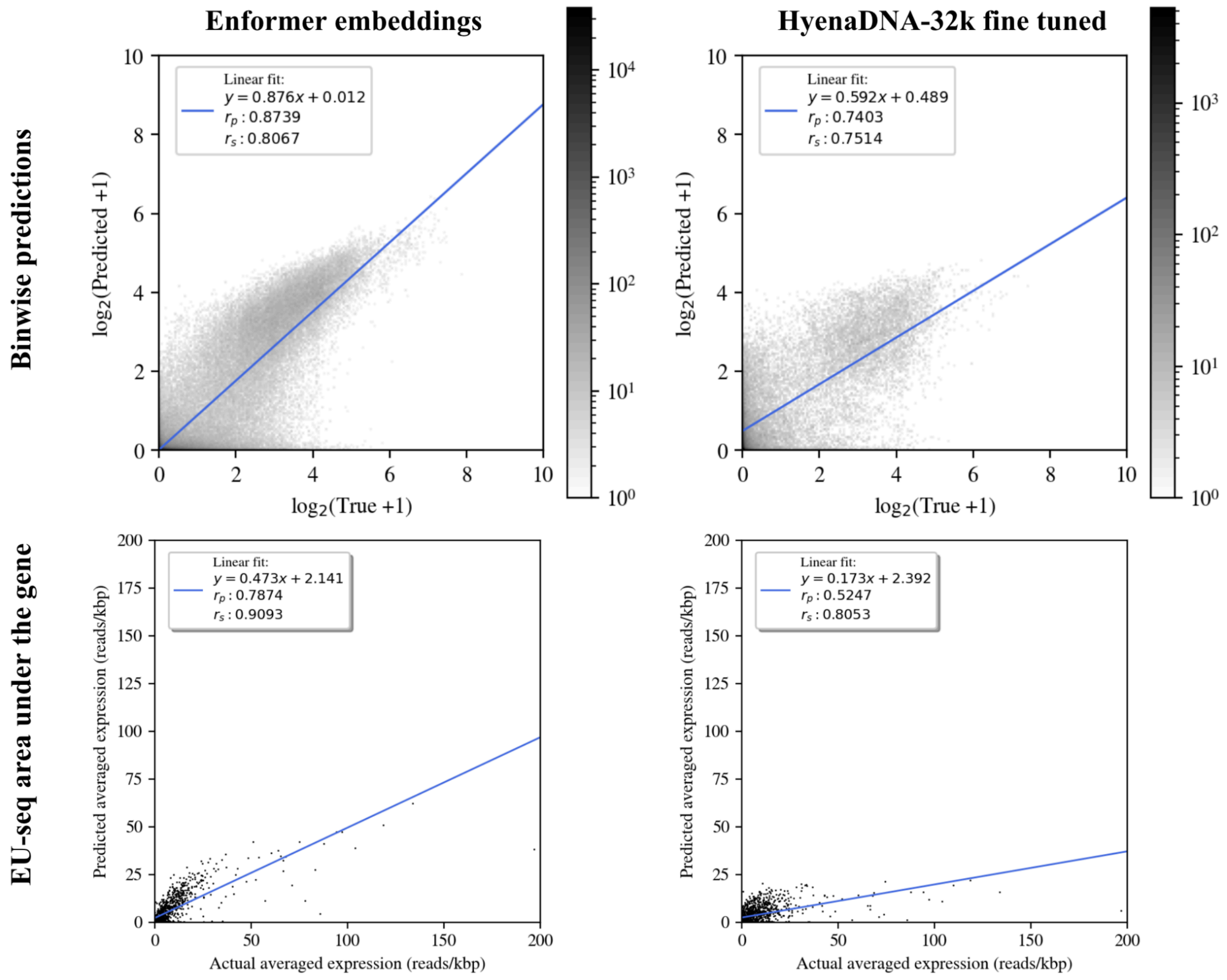


**Supplementary Figure 6. Predicted vs. True target values for benchmark model heads. Top row**) predicted vs. true target values at a bin level, i.e. for every target node describing EU-seq enrichment at a kbp resolution. HyenaDNA-32k predictions are less dense since the narrowed output prediction counted with less targets. **Bottom row)** Predicted vs. True length normalized area under the EU-seq curve inside the reference gene's boundaries. Despite achieving a significant correlation with true targets, HyenaDNA-32k could not reproduce the scale of variation of the EU-seq profiles, which can be seen as a low slope of the fit in the bottom right (m=0.173).

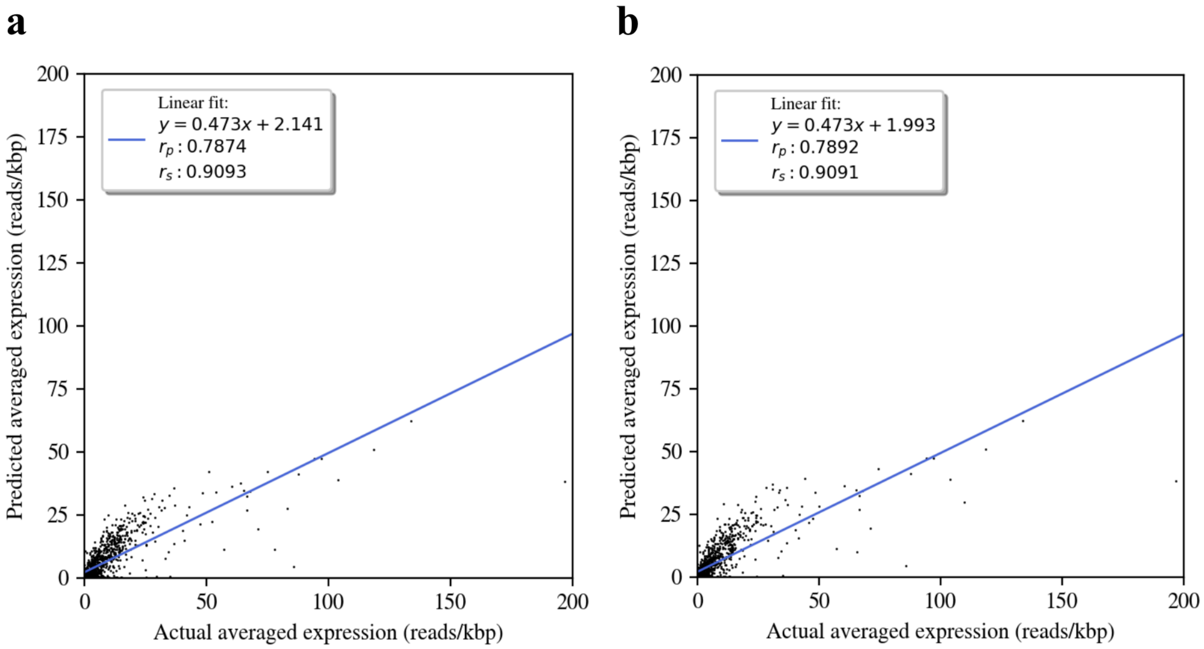


**Supplementary Figure 7. Different target boundaries have little effect on the predicted average EU-seq levels for reference genes.** Reference genes longer than half of the target length were averaged until the boundary. Averaging over the integrated length (length normalization) eases the comparison of different genes. Results shown for Enformer's predictions. **a)** Integrating long genes until Enformer's boundaries (-57kbp, 57 kbp). **b)** Integrating long genes until HyenaDNA-32k boundaries (-15 kbp,+15 kbp).

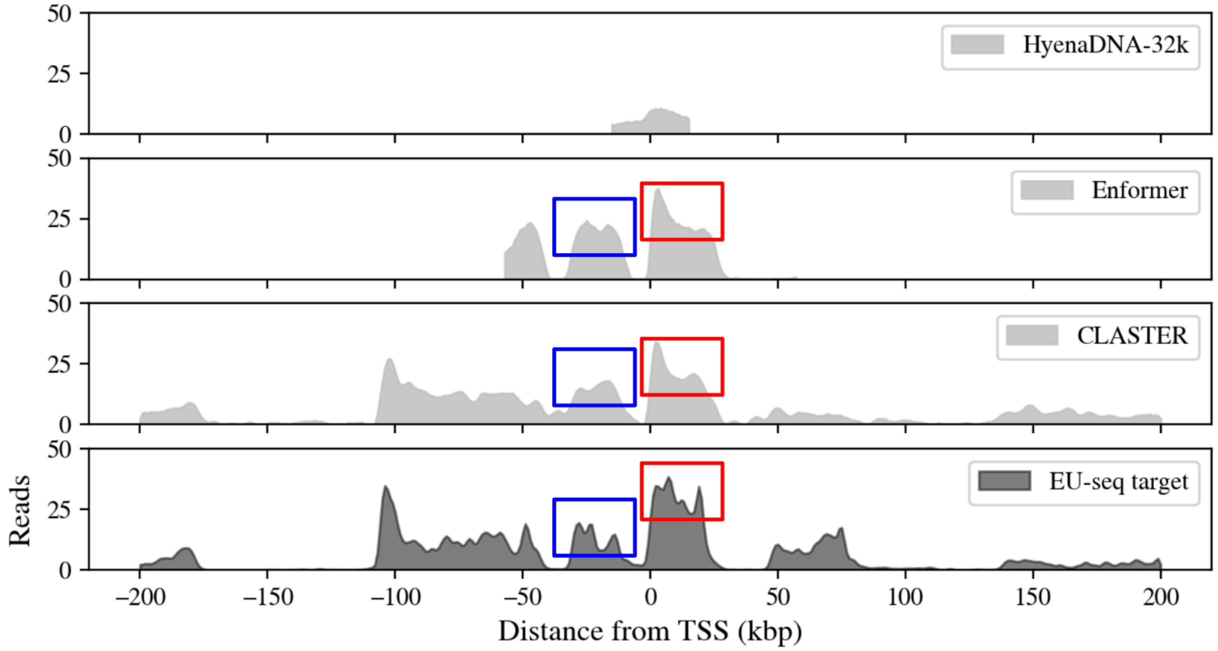


**Supplementary Figure 8. Potential conflicting targets problem might set an upper bound to the accuracy models can achieve.** Profile prediction comparisons between models for samples centered at the gene Nol9 (chr4). Neither the head on top of Enformer embeddings nor CLASTER could fit the high frequency oscillations in the EU-seq targets, suggesting a potential conflicting targets scenario. This occurs when there are multiple possible targets matching a single input, e.g. due to experimental fluctuations in the outputs or the presence of nuanced information in the targets that is not necessarily encoded in the inputs. To minimize the loss, the network's response in such a scenario is to predict the average over potential targets. CLASTER was aimed to predict a wider target window for a given sample to enable us to observe far-ranging changes in output profiles when perturbing single enhancers in the input.

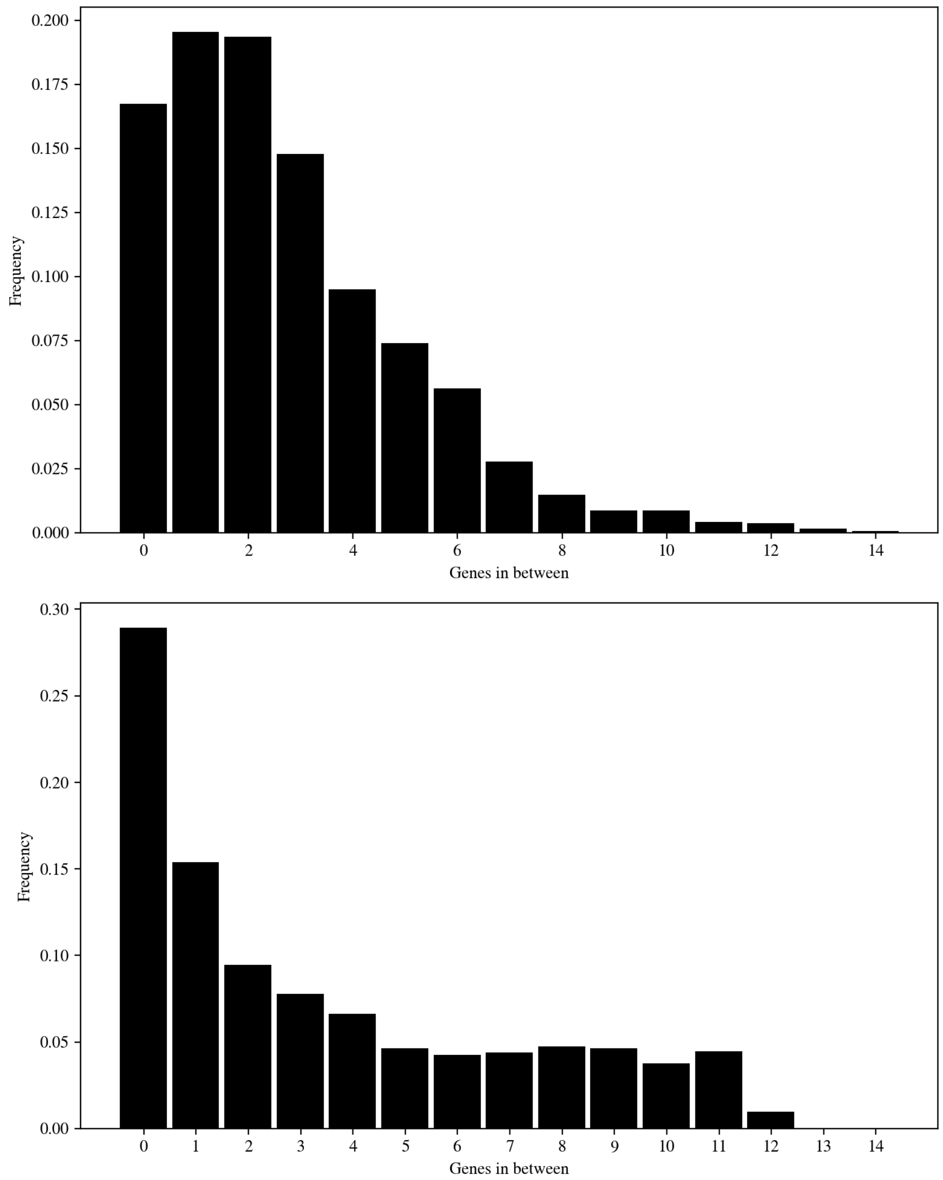


**Supplementary Figure 9. Normalized Enhancer-Gene pair histograms show the same trends.** a) Background histogram counting the number of genes in between each enhancer-gene pair that was kept after data preprocessing, i.e. keeping only active enhancers within a distance of the reference gene in each sample, for the test chromosome (chr4). Enhancer-gene pairs that had few genes in between were more highly represented. b) Histogram keeping only the enhancer-gene pairs for the enhancer that had the highest impact on a gene's expression (histogram in Figure 2) and normalizing by the inverse of the background distribution in a), i.e., if having any number of genes in between had been equiprobable. The low number of occurrences in which a pair had many genes in between boosted the tail of this distribution but still preserved the decay.

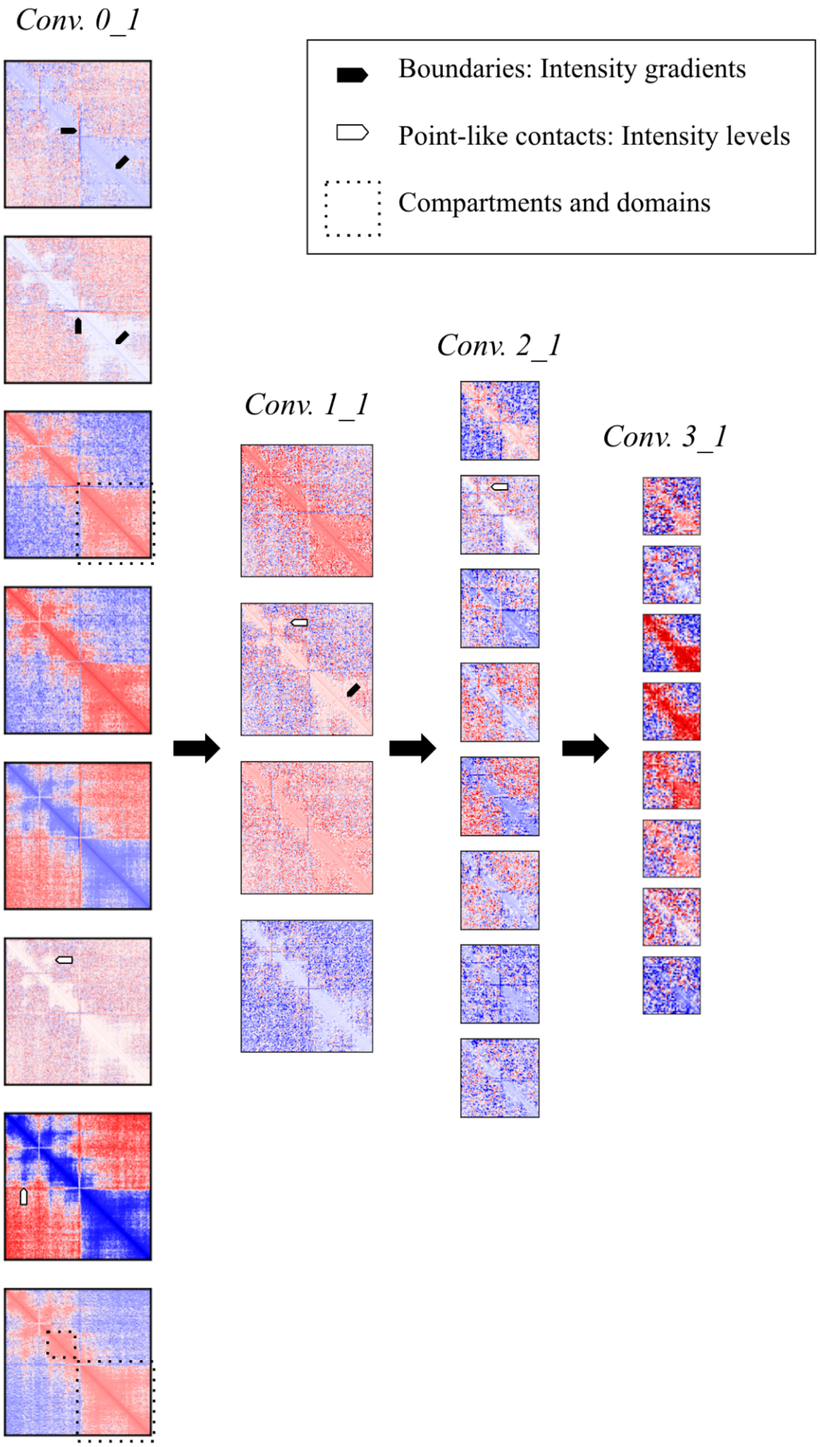


**Supplementary Figure 10. Filter outputs centered at Kif2c (ENSMUSG00000028678.13, chr4)**. The network implemented both high and low pass-like filters, identifying intensity gradients, point-like contacts and high contact compartments and domains. **c)** Deeper filters lose this nuanced information, propagating forward a looser sense of regional contact at wider scales. The symmetry in the data in this setting led to an under usage of the network's capabilities, constraining the size of the final arrays before flattening.

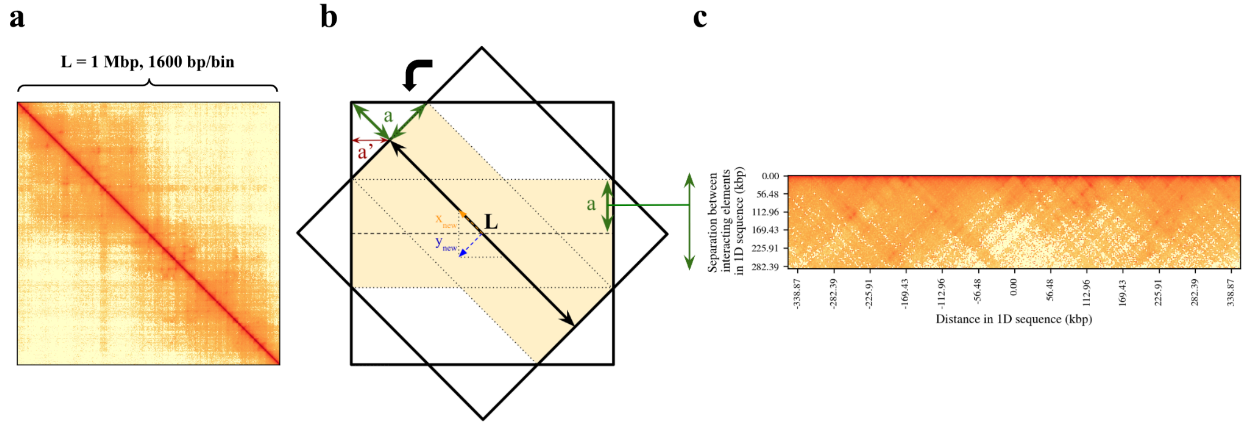


**Supplementary Figure 11. Micro-C matrix rotation and cropping.** Sample centered at the gene Trp53 (chr11). a) Micro-C matrix of shape (625,625), exploring 1Mbp at a 1.6 kbp resolution. b) Rotation and cropping of the matrix. The original squared matrix is rotated 45 degrees and cropped to the maximum possible height to obtain the new rectangular contact map. c) Resulting contact map.

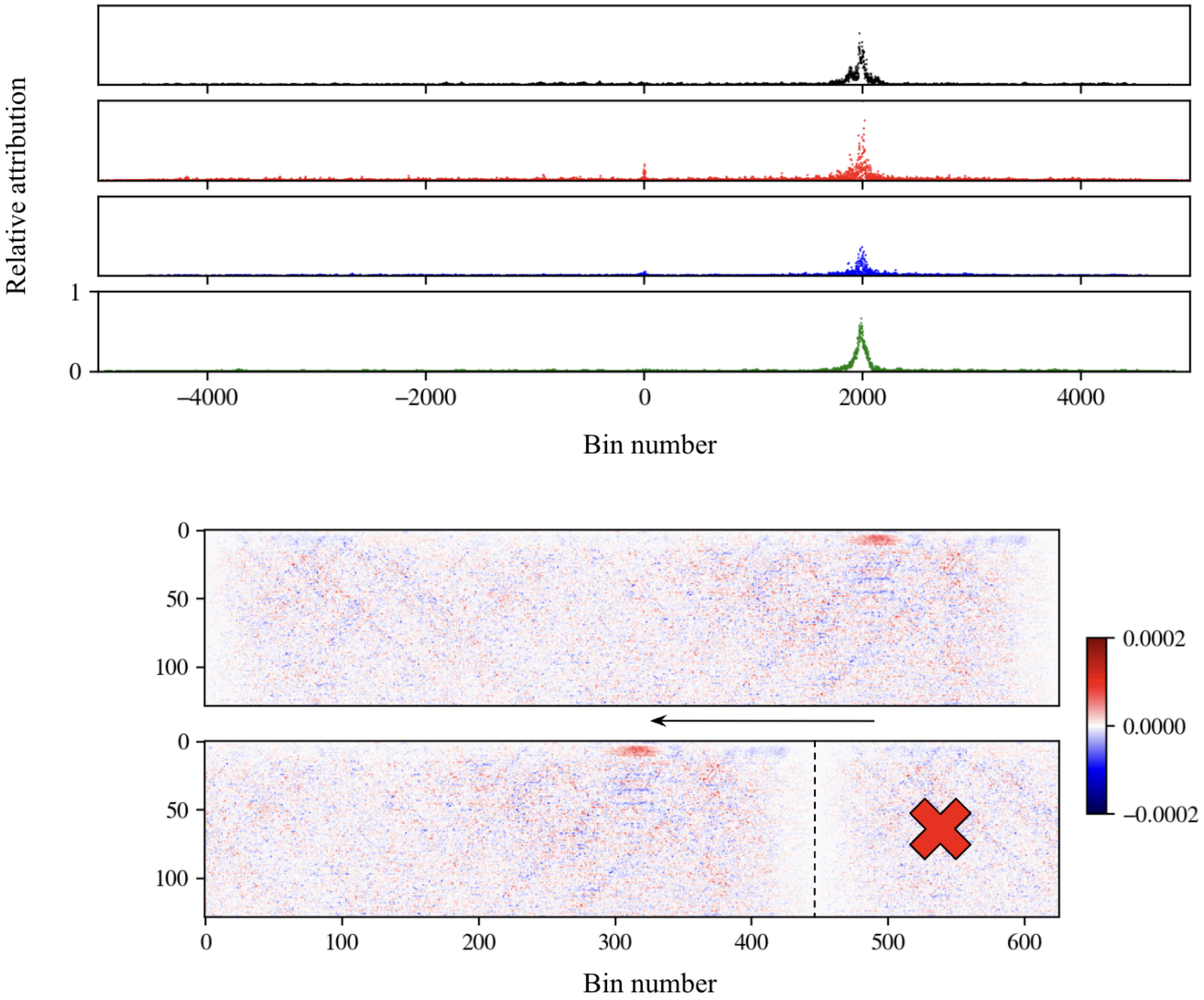


**Supplementary Figure 12. Attributions when predicting expression at +199kbp from the TSS of the central gene. a)** Absolute attributions for chromatin mark tracks relative to the maximum attribution score. **b)** Signed attributions for the rotated and cropped Micro-C map. The final maps were obtained by centering and averaging all maps for different output bins. The original score scale is shown.

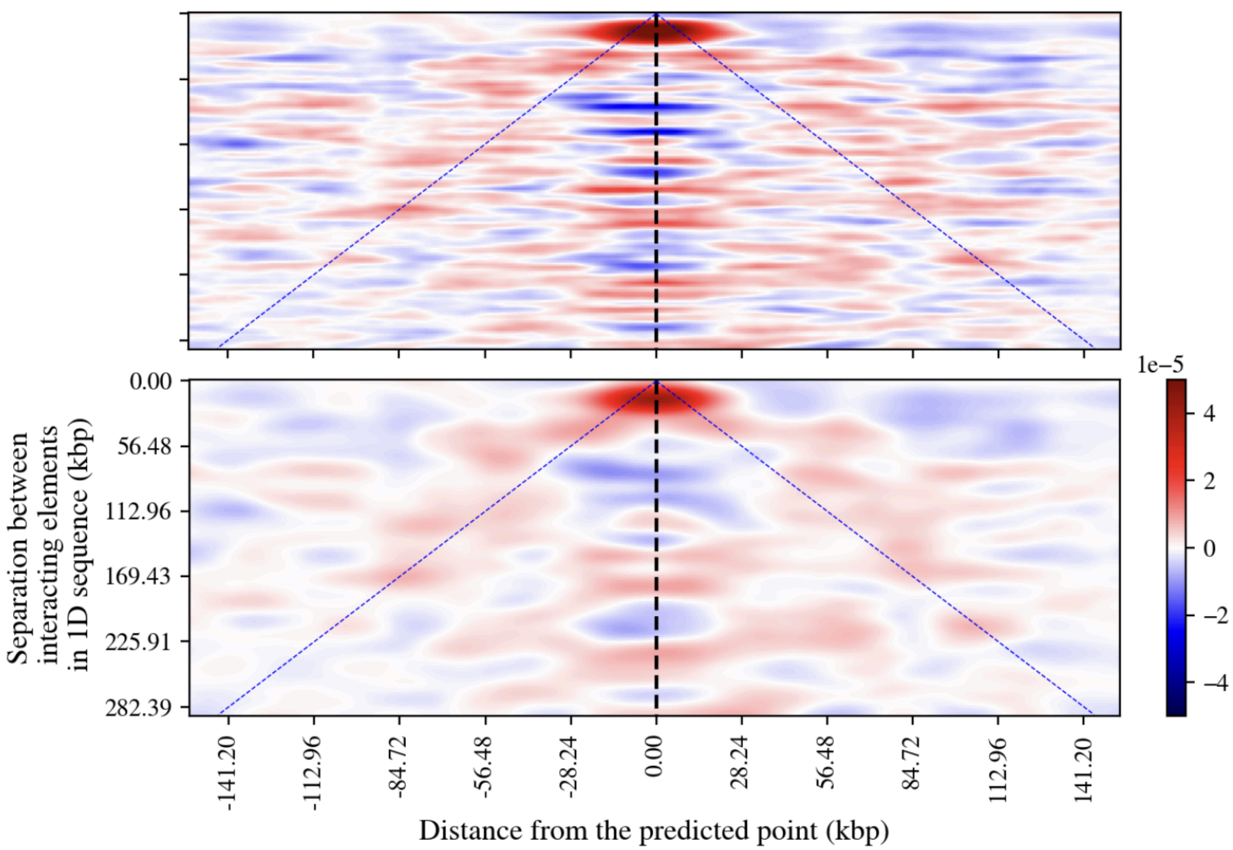


**Supplementary Figure 13. Signed attribution maps smoothed with gaussian filters of varying widths**. Top: attribution score map smoothed with a gaussian kernel of sigma=1. Bottom: attribution score map smoothed with a gaussian kernel of sigma=3. Since the maps corresponding to different predicted points were only scrolled in the horizontal axis, the averages over centered maps yielded a horizontally striped pattern. To compensate for this artifact of the horizontal translations, this gaussian smoothing procedure was applied, allowing us to see more clearly the overall positive attributions associated with regulatory elements in contact with the predicted point. The original score scale is shown.
